# Availability of insects as feed for meadow bird chicks assessed across years by batched image analysis of sticky traps

**DOI:** 10.1101/663591

**Authors:** Charlotte Kaffa, Rutger Vos

## Abstract

As of 1990, there are 27 bird species that have been assigned as meadow birds by the Dutch equivalent of the Farmland Bird Indicator (FBI). These birds have one common characteristic that classifies them as meadow birds: they prefer to breed in meadows. Since 1960, the overall number of meadow birds has been declining rapidly and recently only five species have shown increases. However, not only meadow birds have been declining, this same rate of decline is also seen in many vertebrate, insect, and plant species throughout Europe. Increasing agriculture and urbanisation are considered to be the main causes of these alarming declines and agri-environment schemes show insufficient effect. Not only decreased reproduction rate of meadow birds, but also decreased survival rate of meadow bird chicks may play an important role in the dropping meadow bird numbers. Most of the meadow birds eat insects and it is therefore hypothesized that their food supply is too low. The Louis Bolk Insitute and ANV Water, Land & Dijken have been setting sticky traps in several meadows and counting the number of trapped insects on each sticky trap to assess if the food supply of meadow birds is sufficient. However, counting the insects is very time consuming, unappealing, and error prone. Therefore, a system that uses image analysis to automatically count the insects was improved and deployed as a web application and command line application. This system analyses photographs of sticky traps and counts the insects found on the sticky traps that were set in May 2018. These results were compared to the number of counted insects on the sticky traps that were set in May 2017, tested if the difference was significant and if there was a correlation between the usage of certain management packages. The accuracy of the automated system was also tested by determining if automatically counted results were not significantly different from hand counted results. The results showed that the accuracy of the system was improved but was still unable to provide very reliable results, most likely due to the usage of low-quality photographs from 2017. The number of counted insects from the sticky traps that were set in 2017 was significantly lower as compared to 2018 and no actual correlation could be found between the number of insects and management packages. It is possible for insect populations to have grown this much, however, the difference in insect numbers could have been caused by the difference in temperature when placing the sticky traps, or the sticky traps being less sticky. It is also very likely that the number of insects on the traps in 2017 is lower due to the poor quality of the photographs, so fewer insects could be detected. If the insect populations have grown as significantly as is indicated from the results then it can be stated that the food supply of meadow birds is more sufficient as compared to 2017 and it would be probable that an increase in meadow birds has occurred or will occur in the near future. Further research should be conducted using high quality standardized photographs and carried out for multiple years to gain plentiful reliable data.

## INTRODUCTION

As of 2016, almost one third of the surface area of the Netherlands is made up of meadows (CBS & AgroXpertus, 2018). These fields of grassland are, always or occasionally, favoured for breeding by a group of bird species. This group of birds is appropriately called meadow birds (‘weidevogels’ or ‘boerenlandvogels’), or sometimes also referred to as pasture birds or (wet) grassland birds (Beintema, Moedt, & Ellinger, 1995). The most well-known meadow birds are species that belong to the order of Charadriiformes, such as the Eurasian oystercatcher (‘scholekster’), northern lapwing (‘kievit’), black-tailed godwit (‘grutto’), common redshank (‘tureluur’) and Eurasian curlew (‘wulp’) (Beintema et al., 1995). For most of the meadow birds, as chicks, adults, or both, invertebrates such as insects are their main food source (Beintema & Visser, 1989; ETI Bioinformatics, 2018). Recent studies have shown that there has been a dramatic decline in insects (Hallmann et al., 2017), which raises questions about the sufficiency of food for insect-eating meadow birds.

### Rapid declines

The Dutch equivalent of the Farmland Bird Indicator (FBI), ‘boerenlandvogelindicator’, assigned 27 bird species as meadow birds, table 1. Since 1960, the overall number of meadow birds has declined by 61% up to 73%; 21 meadow bird species have declined in numbers, one meadow bird species remained unchanged in numbers, and only five meadow bird species showed increases. The Eurasian skylark (decline of 96 – 97%), grey partridge (decline of 93 – 95%), European turtle dove (decline of 92 – 95%), Eurasian tree sparrow (decline of 93 – 94%), and godwit (decline of 68 – 79%) show the strongest declines (Boele et al., 2016; Gamero et al., 2017; Gregory et al., 2005; Sovon, 2012). As of 2018, the IUCN Red List (‘rode lijst’), which indicates endangerment based on rarity and negative trends, identifies 12 species of meadow birds that are in danger (Ministerie van Landbouw Natuur en Voedselkwaliteit, 2018).

**Table 1.**
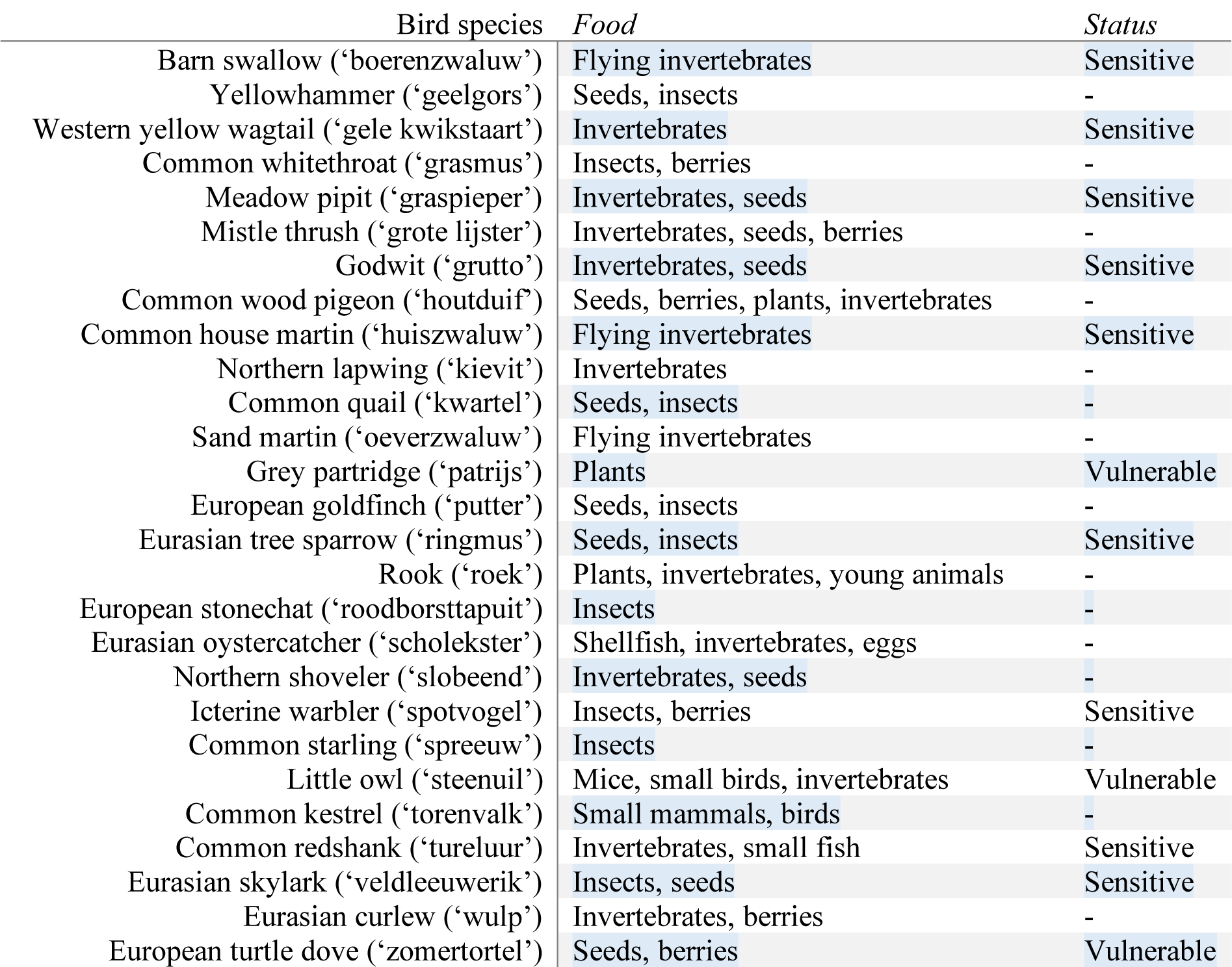
The 27 bird species labelled as meadow birds according to the ‘boerenlandvogelindicator’. Listed for each bird species is their English and Dutch name, their main sources of food, if they are placed on the red list (‘rode lijst’), which indicates endangerment, (‘-’ is stated when the species is not on the red list) and the degree of their endangerment (Ministerie van Landbouw Natuur en Voedselkwaliteit, 2018). Sensitive: stable or increased, but rare, or strong – very strong decline, but still common. Vulnerable: average increase and now quite – very rare, or strong – very strong decrease and now quite rare.

Urbanization and increasingly intensive agriculture are considered to be the main causes of the rapid declines in meadow birds (Chamberlain, Fuller, Bunce, Duckworth, & Shrubb, 2000; P F. Donald, Green, & Heath, 2001; Paul F. Donald, Sanderson, Burfield, & van Bommel, 2006). Fertilization of meadow fields results in higher grass production and earlier mowing, leaving nests unprotected, which leads to increased predation of eggs and chicks. Other agricultural activities such as increased drainage, ploughing, and re-seeding leave soils impenetrable and unsuitable for feeding and breeding (Beintema et al., 1995; Schekkerman, Teunissen, & Oosterveld, 2009; Teunissen, Schekkerman, Willems, & Majoor, 2008).

Most studies show that the declines in meadow bird species are due to decreased reproduction; however, decreased survival rate may also contribute (van Noordwijk & Thomson, 2008), as meadow bird chicks only eat invertebrates, roughly 10.000 a day, that can be found at beak height (Louis Bolk Instituut, 2015). Agri-environment schemes that pay farmers for the provisioning of environmental services effect insufficient change in agricultural activities to restore meadow bird populations (Breeuwer et al., 2009). In addition to postponing mowing dates, reducing fertilization and raising groundwater levels would most likely raise chick survival rates sufficiently (Breeuwer et al., 2009).

Agricultural intensification and specialization does not only affect meadow birds: in Europe, many vertebrate, insect, and plant species have declined at similarly high rates (Benton, Bryant, Cole, & Crick, 2002; Robinson & Sutherland, 2002). Over the last 27 years, average flying insect biomass has declined by 76% up to 82% in protected nature areas in Germany and are considered to have declined as much in the Netherlands. There appears to be no correlation between declines in insects and changing climate. The loss of insect abundance and diversity is expected to lead to cascading effects within ecosystems (Hallmann et al., 2017). Hence, it is hypothesized that the food supply of meadow birds is insufficient due to decreasing insect numbers, causing low survival rate and thus decreasing meadow birds.

### Sticky traps

Louis Bolk Instituut and volunteers from ANV Water Land & Dijken have been working together to assess the food supply of meadow birds in the Netherlands (Louis Bolk Instituut, 2015). Sticky traps, uniform pieces of paper with a sticky surface and bright yellow colour to attract insects, are set up two times in May, and collected after two days. After collecting the traps, the insects trapped on them are then counted by hand. However, this process is very time consuming, error prone, and unappealing. Use of an automated system to count insects will lead to greater insight in insect numbers in meadows, the steps needed to take to halt the decline in meadow bird numbers, and to more possibilities for pursuing large-scale research, as counting the insects is faster, less labour intensive, and free of individual observer biases.

### Aim of research

The aim of the research is to answer the main research question, “Is the food supply for meadow birds in the Netherlands sufficient?” and he subsidiary questions that follow from the main question, “Has the number of insects found on sticky traps that were deployed in the Netherlands declined or increased?”, and “Is there a correlation to be found between insect numbers and the provided management package?”

## METHODS

As prior art for the research, a system that automatically counts insects on sticky traps by image analysis was created, called *sticky-traps*, using the *ImgPheno* library (Naturalis Biodiversity Center, 2018) and tested to determine if its functionality was sufficient (Michels, 2017). *sticky-traps* uses the libraries *OpenCV* 2.4.13, *NumPy* 1.14.5, and *YAML* 1.2. This system was used as a starting point for the web application and command line program, which were used to determine if the food supply for meadow birds in the Netherlands is sufficient.

The *sticky-traps* Python 2.7 script (sticky-traps.py) opens a *YAML* file (sticky-traps.yml) and makes use of the common.py script within the *ImgPheno* library to read the YAML file and return a dictionary object, see figure 1. *YAML*, “YAML Ain’t Markup Language”, (Ben-Kiki & Evans, 2001) is a readable configuration file language which can be easily adjusted without having prior programming knowledge. The *YAML* file is used to provide information about the traps, such as their size, if the edges of the sticky trap need to be cropped and the size of the sticky traps that needs to be cropped. It proceeds to read the images (i.e., photographs of the sticky traps, from here on referred to as “trap”) from the path that was provided, reduces the size of the images, adds the images in a list and converts the colour space of each individual image to HSV using the upper (lightest) and lower (darkest) colour of the trap. This ensures easier detection of the contour of the trap against a dark, preferably black, background.

**Figure 1.**
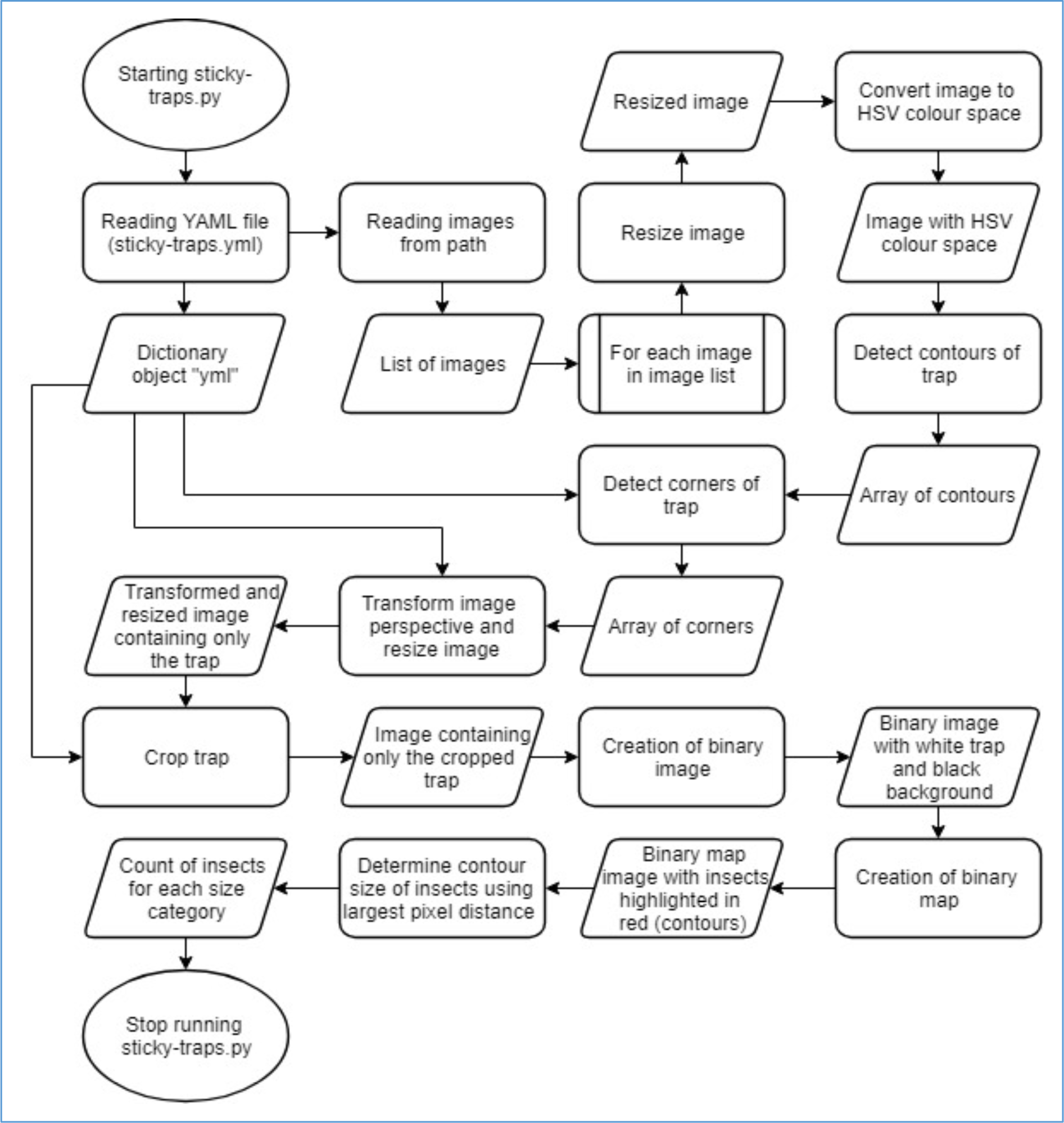
Flowchart of sticky-traps.py. This shows the processes that are carried out when running the sticky-traps.py file, either manually or when called in the web-application, alternating between process and result.

Using the detected contours and the size, provided in *YAML* file, of the trap, it finds the corners of the trap and the image is transformed into a standardised perspective and resized to only fit the trap. The size of the insects can then accurately be determined and the trap is cropped to the dimensions provided in *YAML* file. A binary image is created using the transformed, resized and cropped image; the yellow of the traps is transformed into white and the background into black.

Next, creation of a binary map takes place using the red colour channel, because of its high contrast between the traps and the insects, separating the insects from the trap. Insects can then be segmented into individuals whose contours are detected. The area contained in the detected contours is used to determine the size of the insects by finding the largest distance from one corner to another and counting the pixels in between them and counting the number of insects for each size class (smaller than 4mm, between 4mm and 10mm, larger than 10mm).

### Improvement of the algorithm

The automated system, *sticky-traps*, can induce errors where insects overlap, making it difficult to segment them into individuals; however, in the 2017-version implementation this was the fastest way of running the analysis (Michels, 2017). Therefore, adjustments were made to *sticky-traps* to improve the segmentation of individual insects that lead to a more accurate result in the number of counted insects for each size class. After the sufficiency of each change was tested, the elements of the algorithm that were adapted include different pixel sizes for millimetre classes, adjusted thresholding, changed contour retrieval mode, and deletion of the bilateral filter.

The 2017-version algorithm of sticky-traps uses a slightly altered pixel number for a single millimetre, using 12px as 4mm and 40px as 10mm. This was changed to have cut-off values, no smaller than ∼ 1 millimetre and no larger than ∼ 12 millimetre, which reduces false positives, and a more accurate pixel to millimetre ratio (1mm ≈ 4px), using 15px as 4mm and 38px as 10mm. The adaptive threshold was reduced to a smaller block size of 41 (px), and a larger constant, of 22, that is subtracted from the mean. Furthermore, the contour retrieval mode was changed from the external retrieval mode, retrieving only the outer contours, to the tree retrieval mode, retrieving all the contours and constructing a full hierarchy of nested contours. Last, the bilateral filter that was implemented to eliminate fine texture from the image causes the runtime to be ten times as long and was removed from *sticky-traps*.

Some photographs of traps that were set in previous years could not be analysed by the automated system, as these traps are coloured blue. Analysis of sticky traps with a different colour was implemented by replacing the static colour variables with user input, setting upper (lightest) and lower (darkest) colour of the used traps, from the *YAML* file. *sticky-traps* and the *YAML* file were furthermore adjusted to set if more detailed size classes (0 to 1mm, 1 to 4mm, 4 to 7mm, 7 to 12mm, larger than 12mm) and a result file are preferred. There was an incompatibility issue with the newest version of *OpenCV* (3.4.*) and the automated system would only work on an outdated version, this was solved by changing the output variables of the *OpenCV* findContours-function in *sticky-straps* and the *ImgPheno* source code.

#### Analysis of the algorithm

To test the reliability of the algorithm different methods were tested. The hand counted results of the traps from 2017, 1-5-17 (1^st^ day) and 15-5-17 (2^nd^ day), were used to test various changes of the algorithm by comparing them to the results of automatically counted photographs of the same traps and the deviation was documented. Six methods of detecting insects were tested starting with the algorithm as it was in 2017 (“Automatically counted 101, 10 - w. filter - 1st v. R. Michels”). The six different methods that were tested and their set options can be seen in table 2.

**Table 2.**
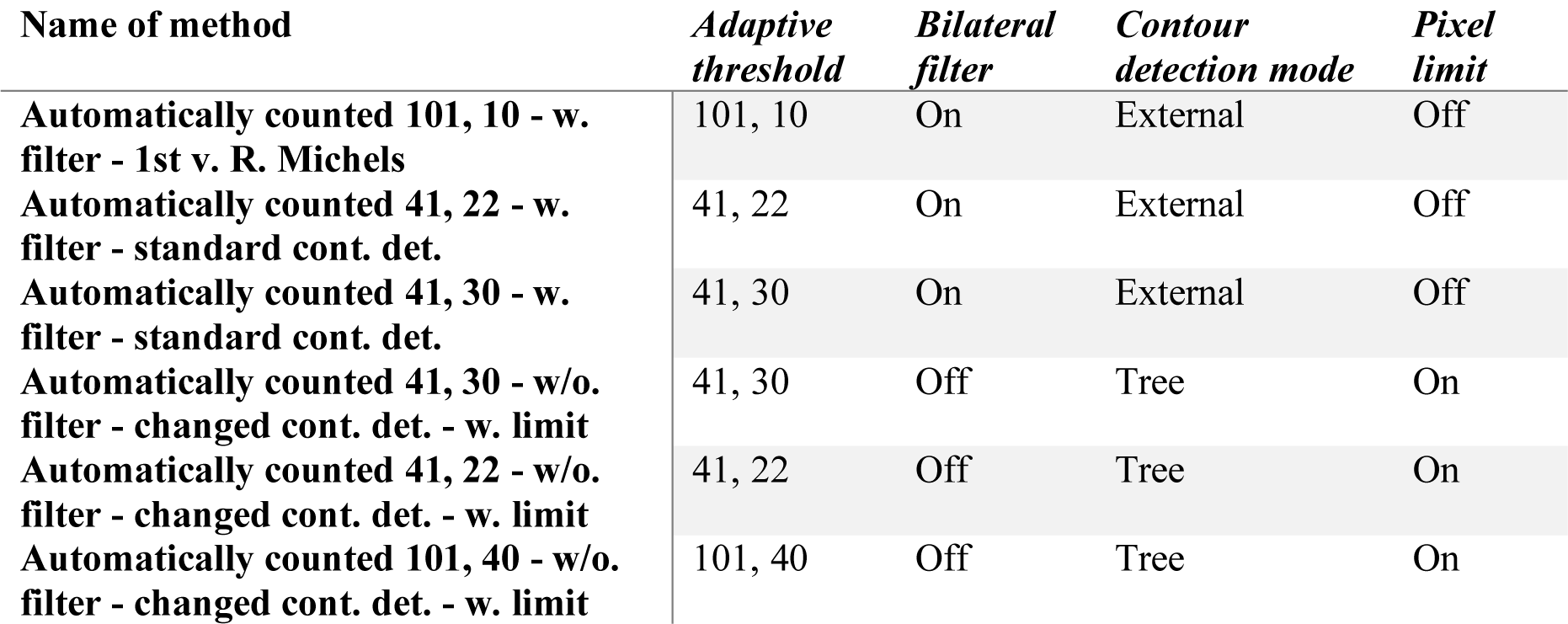
The six methods that were used to test the reliability of the algorithm. The names of the methods contain the different options that were changed to test their sufficiency, automatically counted meaning it was counted by the sticky-traps automated system. The adaptive threshold column specifies the block size and subtracted constant for the adaptive threshold which can identify the insects on the trap. Bilateral filter states if the bilateral filter was implemented, this eliminates noise from the image. If the contour detection was set to retrieve the contours in tree-mode it was set to “Tree”, which constructs a full hierarchy of the retrieved contours, and if it was unchanged it remained “External”, which only retrieves the outer contours. If there was a limited pixel size of 45px, to eliminate extremely large contours, for the larger-than-10 class then “pixel limit” is on.

| **Name of method** | ***Adaptive threshold*** | ***Bilateral filter*** | ***Contour detection mode*** | ***Pixel limit*** |
| --- | --- | --- | --- | --- |
| **Automatically counted 101, 10 - w. filter - 1st v. R. Michels** | 101, 10 | On | External | Off |
| **Automatically counted 41, 22 - w. filter - standard cont. det.** | 41, 22 | On | External | Off |
| **Automatically counted 41, 30 - w. filter - standard cont. det.** | 41, 30 | On | External | Off |
| **Automatically counted 41, 30 - w/o. filter - changed cont. det. - w. limit** | 41, 30 | Off | Tree | On |
| **Automatically counted 41, 22 - w/o. filter - changed cont. det. - w. limit** | 41, 22 | Off | Tree | On |
| **Automatically counted 101, 40 - w/o. filter - changed cont. det. - w. limit** | 101, 40 | Off | Tree | On |

The six methods were tested on photographs of the traps that were made with a professional camera, while the traps were in a protective plastic cover, and on photographs made by the volunteers, while some traps were in a protective plastic cover. For each method the number of exact matches, results within 10% deviation, results within 20% deviation, and results with more than 20% deviation were counted. Using 0-10% deviation as an acceptable deviation or correctly counted results and >10% deviation as incorrectly counted results. From these results an average deviation for each trap was calculated and an absolute deviation for each trap was calculated. To compare the methods all the average deviation results and absolute deviation results were used as an average-average deviation and an average absolute deviation per method. The method with the highest average-average 0-10% deviation, lowest average-average >10% deviation and lowest average absolute deviation was the best performing method and was then chosen to perform the analysis of all the traps. The same method was also applied to show the difference in results for a trap when a different photograph was used (photographs made with professional camera versus photographs made by the volunteers). Next, the significance for the difference between the hand counted results and automatically counted results was calculated, using the method used in 2017 and the method that was chosen as the best performing method, for both days in 2017 and 2018 and using photographs taken with a professional camera and photographs taken by the volunteers. This was calculated using two-sided paired t-tests, for all size classes, to make sure that the difference between the hand counted results and automatically counted results were not significantly different and therefore more accurate.

#### Performance of the automated system

Five datasets were formed randomly containing various photographs of traps from 2017 and 2018, with images of different sizes. Starting from dataset “1”, as the smallest dataset with the least files and the smallest size in MBs (megabytes), up to dataset “5”, the biggest dataset with the most files and the largest size in MBs, as seen in table 3. The runtime for each dataset was then documented for the method that was used in 2017 and the method that was tested as the best performing method. These results were plotted in a graph and a trend line was added.

**Table 3.** Overview of the datasets that were used for testing the performance of the automated system. The table contains the number of the dataset from 1 to 5, the number of files in the dataset and the size of the dataset in MBs. Also, a total overview of the number of files and the size in MBs.

| ***DATASET*** | ***FILES*** | ***SIZE*** |
| --- | --- | --- |
| ***1*** | 80 | 202 MB |
| ***2*** | 147 | 231 MB |
| ***3*** | 281 | 605 MB |
| ***4*** | 549 | 1360 MB |
| ***5*** | 1085 | 2960 MB |
| ***TOTAL*** | **2142** | **5360 MB** |

#### Web application

A web application was set up to aid in user friendly uploading of photographs of traps, providing analysation of the traps with the automated system *sticky-traps.* A cloud instance was provided and *Apache HTTP Server* 2.4.33, *Django* 1.11, *SQLite* 3.24.0, and *Geoposition* 0.3.0 were installed. This web application was based on the *OrchID* (Pereira, Gravendeel, Wijntjes, & Vos, 2016) web application that uses the *NBClassify* library (Naturalis Biodiversity Center, 2017). The web application was based on a Model-View-Controller model, as seen in figure 2. The user is able to submit data as input at the controller level, this data manipulates the model level and it creates a relationship between the data and the created models. The data and requests are transmitted to the view level, this updates the view level. At the view level the data is saved into the database and, in case of the *sticky-traps* web application, the process of analysing the data is carried out. When this process has completed the view level will respond with, in the case of the *sticky-traps* web application, a result page. The user is then able to see this result page.

**Figure 2.**
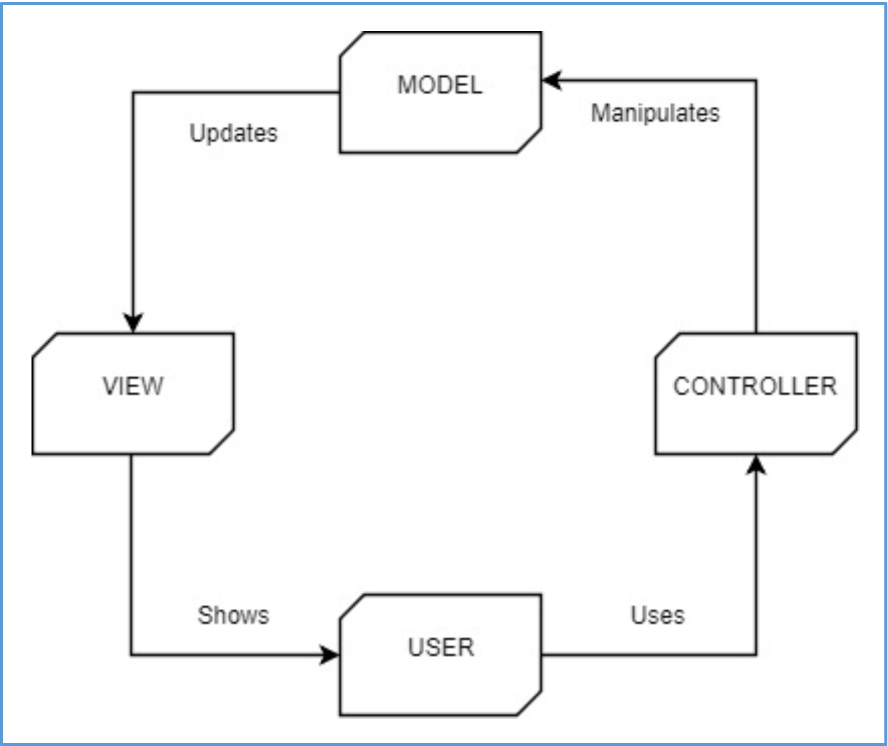
Model-View-Controller model overview. Django is based on a Model-View-Controller model. It allows users to provide input at the controller level, the controller level then manipulates the model and creates relations between the input and the created models. The data and request are transmitted to the view level, this updates the view level and displays a response that can be seen by the user.

*Django* makes use of this Model-View-Controller model and it consists of several Python library files that work together. The files that were set up consist of a settings file, an admin file, a forms file, a models file, an URLS (Uniform Resource Locator) file, and a views file. The *settings* file contains all the settings to correctly run the web application via *Django*, such as the base directory, the allowed hosts, installed applications that are needed to run the web application, the database, and *Geoposition* options. The *admin* file is used to manage the internal organisation of the web application. Using *forms* ensures that the web application can receive user input, it holds the classes for the models that are created in *models*, which defines all the fields for each model. Specifying specific URL patterns in *urls* creates URLS that, e.g., do not contain a file format. Finally, *views* takes a request and returns a response. In the *sticky-traps* web application *views* is requested to display a homepage, an upload page and saving the user input data into the database, analysing the data by calling *sticky-traps* and displaying a result page. To display the homepage, upload page and result page *views* uses the *HTML* files *homepage, upload, results* and *base_layout* to display them. The *HTML* files make use of the installed *CSS* and *JS* files which contain the information for the format and design of the *HTML* pages. After the setup of the *Django* files was completed a subdomain was created, called “plakvallen.naturalis.nl”, and the settings were changed to display the web application on this subdomain.

#### Command line program

Due to increased interest of other research facilities, a command line version of *sticky-traps* was created in addition to the web application. The command line version was specifically made for large-scale research and is suitable to use for the analysis of large amounts of photographs of traps. The command line version can be easily adjusted to fit the traps that have been used in fieldwork by adjusting the *YAML* file, as mentioned on page 5 and 7. The automated system is part of the *ImgPheno* library and can be found in the examples. After installation of *ImgPheno* is complete the automated system can be executed by entering “python sticky-traps.py” in a terminal window. It will analyse all the images that are found in the folder “imgpheno/examples/images/sticky-traps”. A few changes were made to *sticky-traps* for it to run on command line, as it no longer receives a request from the *sticky-traps* web application. These changes include specification of a direct path to the images, creation of an output file and output that is printed on the command line, implemented detailed size classes and changeable trap colour.

### Fieldwork

The traps were deployed on two days, 7-5-18 (1^st^ day) and 28-5-18 (2^nd^ day), by the volunteers of ANV Water, Land & Dijken in 19 fields with various management packages. The management packages, indicating which management style was used on a field, are listed in table 4. The companies the traps were placed on are Van der Deure (LB, UM), Engewormer (VH, WP), Van de Hudding (EW, KG, LB), Koning (KG, LB, UM), Vlaar (KG, LB), and Westerneng (LB, UM). To achieve accurate results the traps that were used to capture the insects were uniform in size, 10cm × 24.7cm, and in colour, yellow without printed lines. In each meadow the traps were positioned in a diagonal line with gaps of 10 meters between each trap and the edge of the field. Figure 3 shows a photograph and diagram of how the traps were positioned in the field. To avoid getting grass or soil on the traps the grass around the traps was trimmed and the traps were placed about 15cm above the ground vertically. Two days after placement the traps were collected from the meadows and photographed by the volunteers.

**Table 4.** The management packages used by the companies to indicate the management style that was used on a field. The table consists of the abbreviation and the English meaning for each abbreviation, the Dutch meanings are listed in brackets.

| ***Abbreviation*** | ***Meaning*** |
| --- | --- |
| ***EW*** | Extensive grazing ('Extensief weiden') |
| ***KG*** | Herb-rich grassland ('Kruidenrijk grasland') |
| ***LB*** | Nest management ('Legselbeheer') |
| ***UM*** | Postponed mowing ('Uitgesteld maaien') |
| ***VH*** | Moist meadow ('Vochtig hooiland') |
| ***WP*** | Meadow bird package ('Weidevogel pakket') |

**Figure 3.**
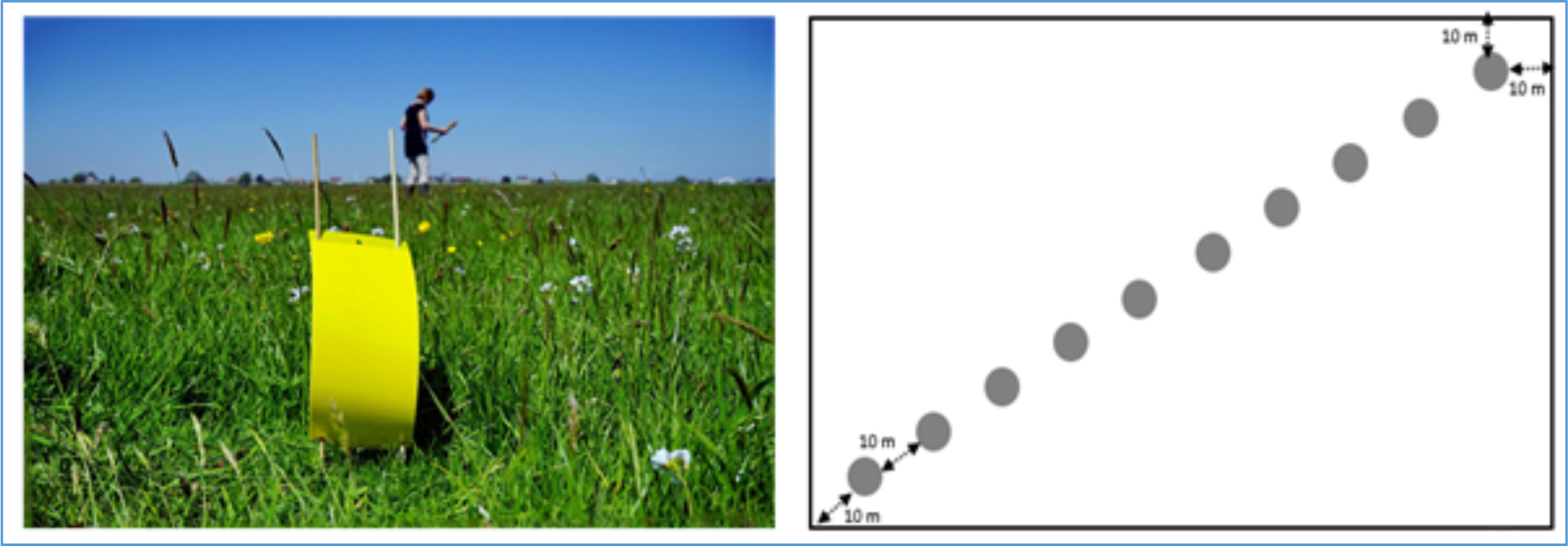
Positioning of the traps. On the left a deployed trap is shown (ANV Water Land & Dijken, 2018), it is placed vertically and about 15cm above ground level. Without any insects trapped on the trap it can be seen that the trap has no printed lines and has a uniform yellow colour. On the right a visualisation is shown of the placement of the traps on the field. There ten traps are deployed on each field with ten metres between each trap and the sides of the field.

### Analysis of the traps

To assess the number of insects found on the traps and whether an increase or decline has occurred, all the traps, both days in 2017 and both days in 2018, were analysed using the method that was chosen as the best performing method.

#### Calculating significance

The results of the analysis of the traps were plotted in bell curves, using the mean and standard deviation, to visualise if the data was normally distributed. From the counted total insects, smaller than 4mm, between 4 and 10mm, and larger than 10mm averages were formed for each meadow. Next significant difference was calculated for all meadows, for all classes (total, <4mm, 4-10mm, >10mm) on both days in 2017 and 2018 by performing paired two-sided t-tests for averages. To calculate significant difference between each individual meadow on both days in 2017 and 2018 for all classes the raw counts were first tested with f-tests to assess if the variance was significantly different; next, two-sided t-tests with equal or unequal variances (depending on the result of the f-test) were performed. For the average total counts for each individual meadow in 2017 and in 2018 the averages were grouped per management package and a single factor ANOVA was performed to test significant difference between management packages in one year.

## RESULTS

### Results and significance of the algorithm

Not all of the traps from 2017 (various photographs made with a professional camera, “prof”, and some made by the volunteers, “vol”) are hand counted; a few from Van der Deure (LB, UM), Engewormer (VH2, VH3), Van de Hudding (EW), Koning (KG), Vlaar (KG, LB, UM), and Westerneng (UM) that were set on the 1^st^ day. Also a few of the traps that were set on the 2^nd^ day are hand counted; Van der Deure (UM), Van de Hudding (EW, KG), Koning (KG, LB), and Westerneng (UM). For all the traps that were hand counted the prof images are available and are analysed and a smaller number of vol images are available and analysed as well. Figure 4 shows a part of the overview of these results and the absolute and percentage of deviation, the complete file can be found in the appendix. An overview of the comparison of these results to the hand counted results for each method can be seen in figure 5.

**Figure 4.**
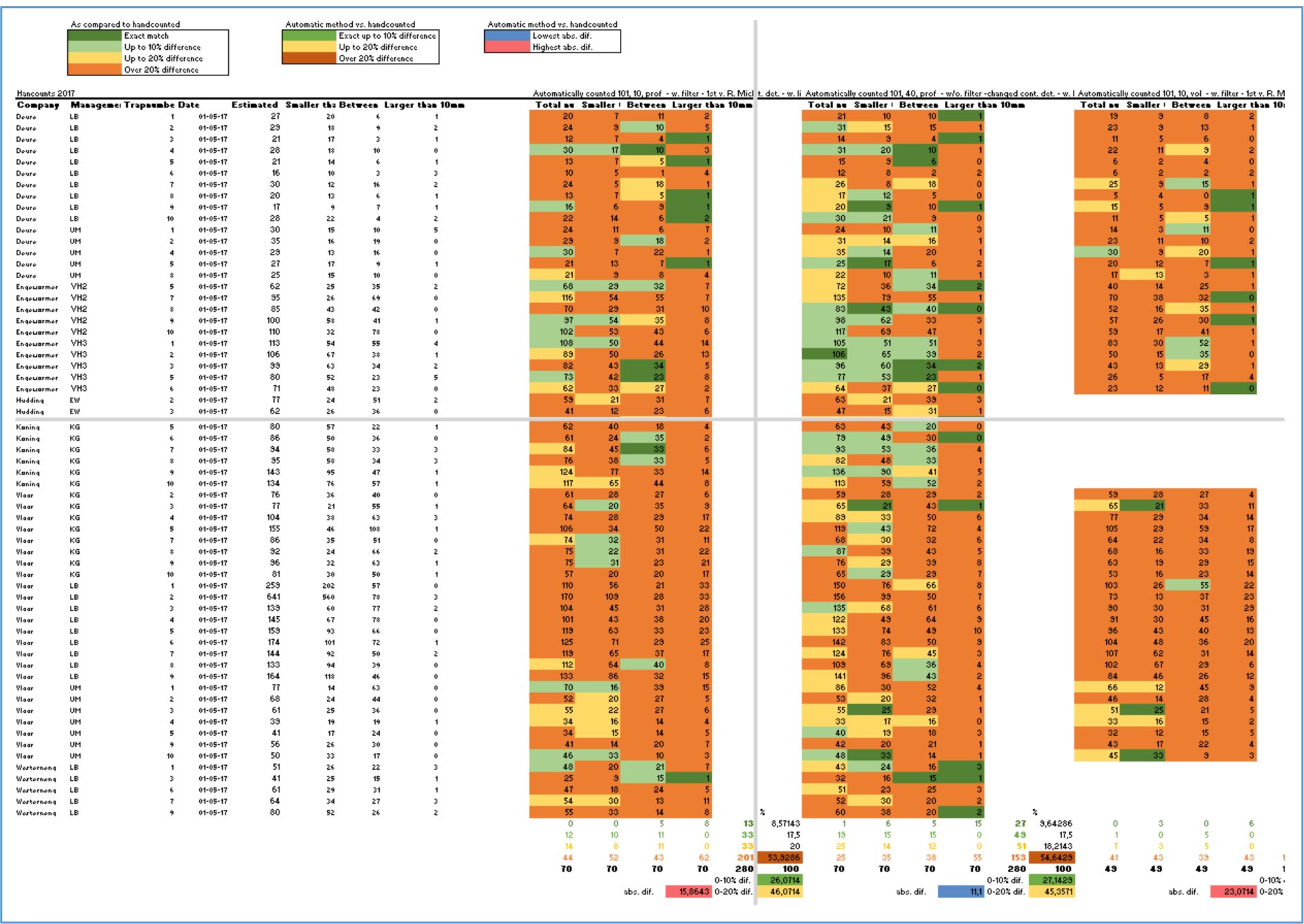
Part of tab “Accuracy Automated System” from the file “Sticky-traps results 2018 and overview”. This tab shows results for the different algorithm methods that are used, comparing them to the hand counted results, to show their sufficiency. All traps that are used for this comparison are from 2017. On the left are the company names, management packages, trap numbers, and the date of deployment of each trap. Next are the hand counted results, starting with a total estimate, the number of insects that are smaller than 4mm, between 4 and 10mm, and larger than 10mm. Thereafter are the automatically counted results, with the same size classes, for the tested methods for the algorithm; “Automatically counted 101, 10 - w. filter - 1st v. R. Michels”, “Automatically counted 41, 22 - w. filter - standard cont. det.”, “Automatically counted 41, 30 - w. filter - standard cont. det.”, “Automatically counted 41, 30 - w/o. filter - changed cont. det. - w. limit”, “Automatically counted 41, 22 - w/o. filter - changed cont. det. - w. limit”, and “Automatically counted 101, 40 - w/o. filter - changed cont. det. - w. limit”. These methods are tested for the prof images and vol images, for both days in 2017. On the top it shows three legends, one corresponding to the percentage of deviation for each individual result as compared to the hand counted traps (dark green = exact match, light green = up to 10% difference, yellow = up to 20% difference, orange = over 20% difference), one corresponding to the average absolute deviation per trap as compared to the hand counted traps (blue = lowest absolute difference, red = highest absolute difference), and one corresponding to the average percentage of deviation per trap as compared to the hand counted traps (green = exact up to 10% difference, yellow = up to 20% difference, red = over 20% difference). Below each method is an overview of the total frequency for exact matches, up to 10% difference, up to 20% difference, and more than 20% difference, and an average of these categories for each trap. It also shows the average absolute difference for each trap. The first two colour-coded tables are both prof images, the third are vol images, here can be seen that a number of vol images are missing. The results in this figure are presented in a Dutch format and therefore a comma is used instead of a dot to display decimals.

**Figure 5.**
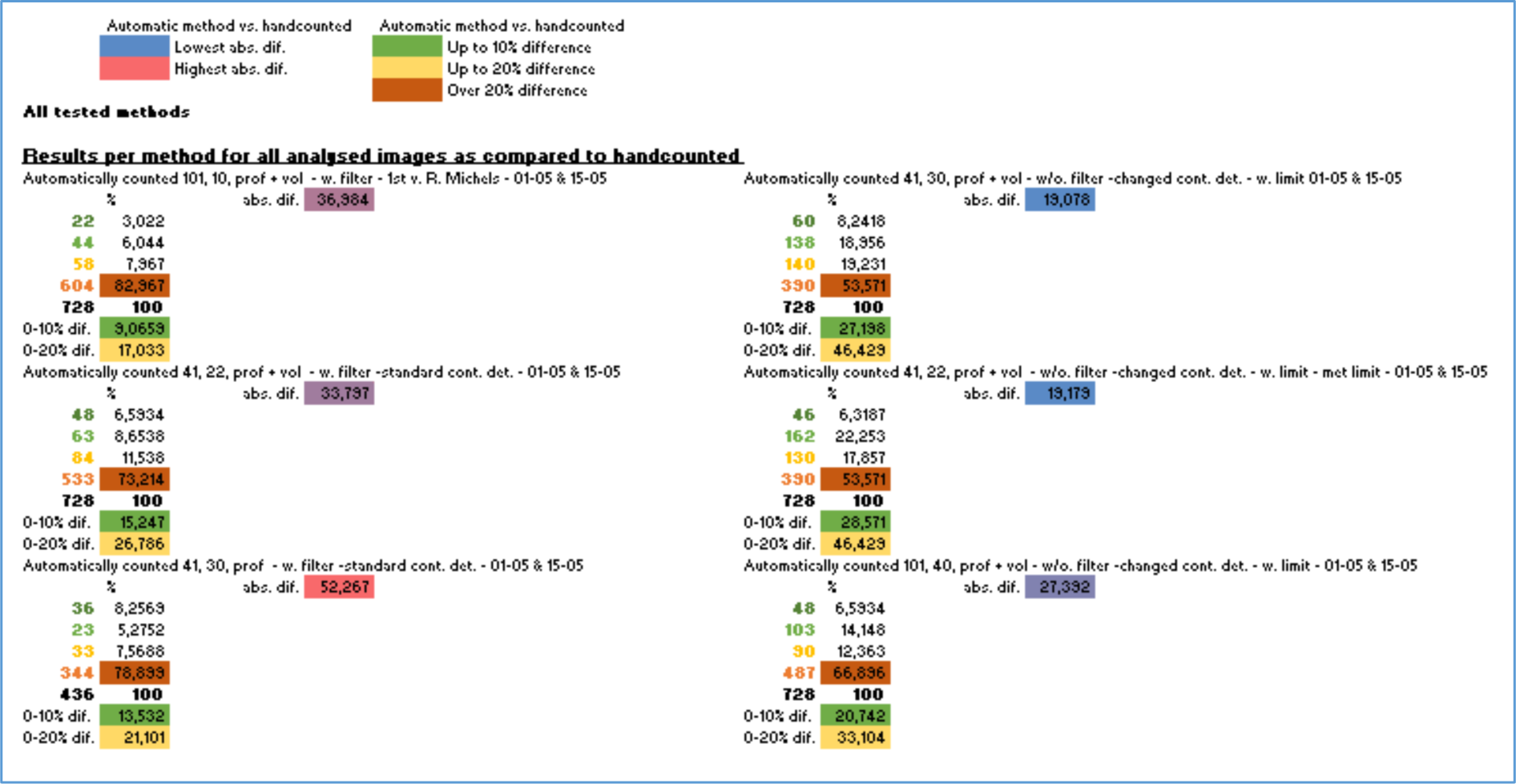
Part of tab “Summary Methods” from the file “Sticky-traps results 2018 and overview”. This tab shows the average results for each of the methods for the algorithm that were used to test their sufficiency. These methods are tested for the prof images and vol images, for both days in 2017. At the top it shows two legends, one corresponding to the average absolute deviation per trap per method as compared to the hand counted traps (blue = lowest absolute difference, red = highest absolute difference), and one corresponding to the average percentage of deviation per trap per method as compared to the hand counted traps (green = exact up to 10% difference, yellow = up to 20% difference, red = over 20% difference). A complete frequency for exact matches, up to 10% difference, up to 20% difference, and more than 20% difference for each method and an average per method is calculated. Also, an average absolute difference for each method is calculated. The results in this figure are presented in a Dutch format and therefore a comma is used instead of a dot to display decimals.

The method “Automatically counted 41, 22 - w/o. filter - changed cont. det. - w. limit” is selected as best performing method, with an average absolute difference per trap of 19.179 and an average deviation of 0-10% for 28.571% of the traps. It has the lowest average percentage of traps that have over 20% deviation. The method that was implemented in 2017, “Automatically counted 101, 10 - w. filter - 1st v. R. Michels”, shows one of the worst results. This method has a very high average percentage of traps that have a deviation over 20%, 82.967%, which are considered as incorrect values, and a large average absolute difference per trap of 36.984.

Bell curves are created to visualise if the data, hand counted total insect counts from 2017 from both days and automatically counted total insect counts from 2017 from both days with the method implemented in 2017 and the best performing method, is normally distributed. These bell curves can be seen in figure 6. The bell curves indicate that the data is normally distributed and a t-test is able to be performed on the data.

**Figure 6.**
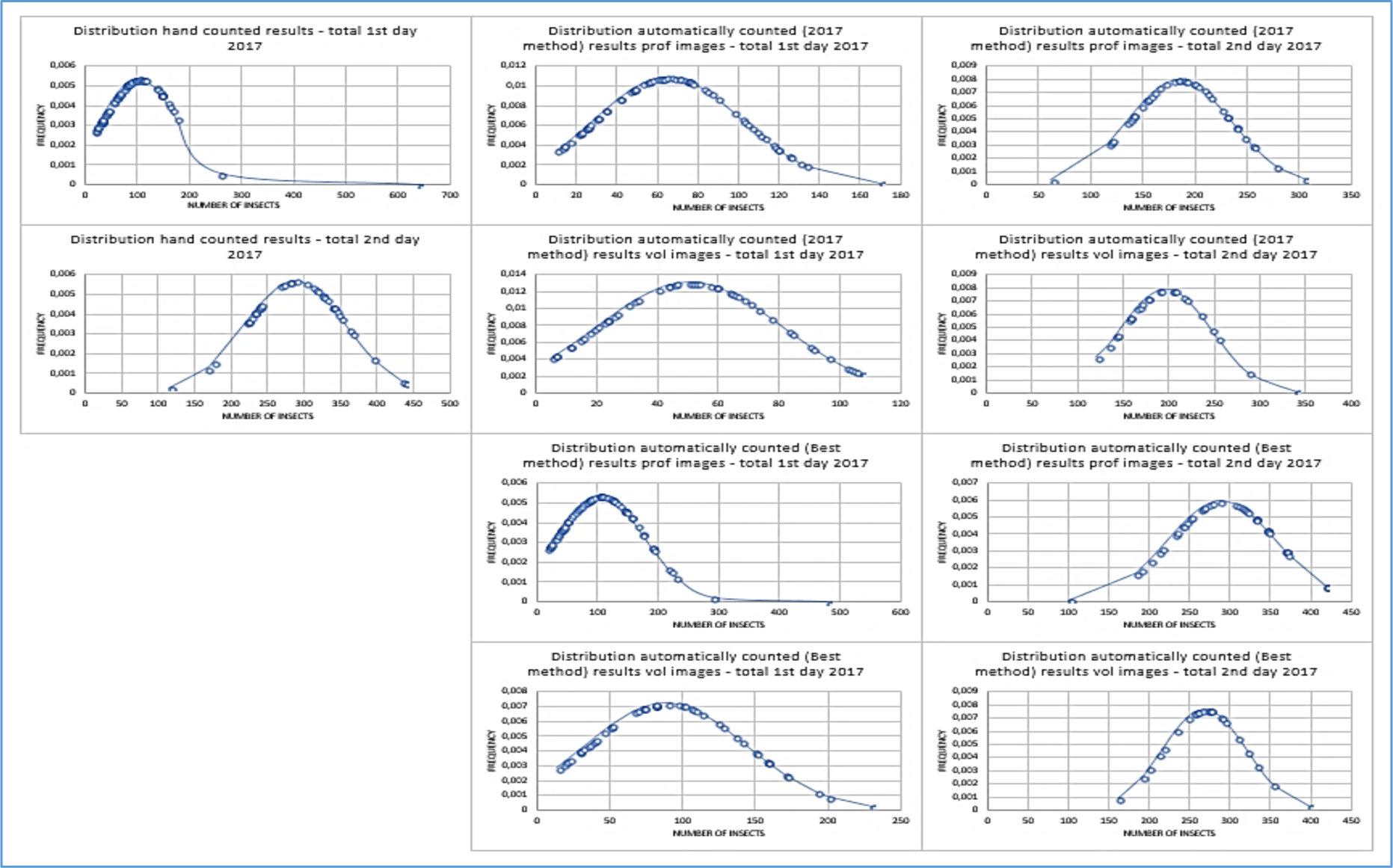
Histograms showing distributions of 2017-method and best method. Leftmost curves visualise the hand counted total insect counts for the 1^st^ and 2^nd^ day. Immediately next are the curves that visualise the automatically counted total insects, using the 2017-method, for the prof and vol images for both days. Beneath those are the curves that visualise the automatically counted total insects, using the best performing method, for the prof and vol images for both days.

The results of the performed t-tests indicate that the best working method more frequently shows that there is no significant difference between the hand counted results than the 2017-method, however a little more than half of the results do show a significant difference. Table 5 contains an overview of these results.

**Table 5.** Two-sided paired t-test results of hand counted and automatically counted results from both days of 2017. The table contains the t-test results for different classes for prof and vol images, alternating between the 2017-method and the best performing method. Significant (α = 0.05) differences indicated with asterisks *.

| Category (vs. hand counted) | 2017-method prof | Best method prof | 2017-method vol | Best method vol |
| --- | --- | --- | --- | --- |
| Total 1<sup>st</sup> day | 0.002* | $2.803 \times 10^{-05}$* | 0.001* | 0.851 |
| Total 2<sup>nd</sup> day | $8.985 \times 10^{-19}$* | 0.756 | $2.225 \times 10^{-08}$* | 0.966 |
| <4mm 1<sup>st</sup> day | 0.007* | $4.360 \times 10^{-04}$* | $8.909 \times 10^{-03}$* | 0.971 |
| <4mm 2<sup>nd</sup> day | $1.650 \times 10^{-17}$* | 0.152 | $3.3108 \times 10^{-07}$* | 0.426 |
| 4-10mm 1<sup>st</sup> day | $3.996 \times 10^{-08}$* | 0.265 | $2.0173 \times 10^{-08}$* | 0.007* |
| 4-10mm 2<sup>nd</sup> day | 0.339 | $3.312 \times 10^{-07}$* | 0.981 | 0.0544* |
| >10mm 1<sup>st</sup> day | $2.154 \times 10^{-11}$* | $4.214 \times 10^{-05}$* | $1.012 \times 10^{-05}$* | $4.417 \times 10^{-05}$* |
| >10mm 2<sup>nd</sup> day | $9.906 \times 10^{-15}$* | 0.002* | $3.572 \times 10^{-09}$* | 0.0115* |

Most results of the performed t-tests, as seen in table 6, to indicate if there is significant difference between the prof and vol images using the 2017-method and the best performing method show that there is significant difference between them. The best performing method does not have significantly less difference between the prof and vol images.

**Table 6.** Two-sided paired t-test results of automatically counted results from both days of 2017 of prof versus vol images. Significant (α = 0.05) differences indicated with asterisks *.

| Category | 2017-method prof vs. vol | Best method prof vs. vol |
| --- | --- | --- |
| Total 1<sup>st</sup> day | $2.728 \times 10^{-06}$ * | $1.984 \times 10^{-03}$ * |
| Total 2<sup>nd</sup> day | $3.006 \times 10^{-06}$ * | $2.604 \times 10^{-05}$ * |
| <4mm 1<sup>st</sup> day | $3.685 \times 10^{-05}$ * | $9.757 \times 10^{-03}$ * |
| <4mm 1<sup>st</sup> day | 0.019 * | $8.346 \times 10^{-04}$ * |
| 4-10mm 1<sup>st</sup> day | 0.859 | $9.582 \times 10^{-06}$ * |
| 4-10mm 2<sup>nd</sup> day | $6.582 \times 10^{-04}$ * | 0.689 |
| >10mm 1<sup>st</sup> day | $1.283 \times 10^{-08}$ * | 0.073 |
| >10mm 2<sup>nd</sup> day | 0.759 | $5.624 \times 10^{-05}$ * |

#### Performance of the algorithm

After analysis of the five different sized datasets the best performing method ran almost twice as fast with a smaller sized dataset (dataset 1, 2) and a little slower with a larger dataset (dataset 3, 4, 5), as compared to the 2017-method, as can be seen in table 7. Both methods have a big O of O(2^n^). The fitted trendline of the 2017-method is y = 42.167e^0,7767x^ and the fitted trendline of the best performing method is y = 19.077e^0,9621x^, this is visualised in a graph in figure 7.

**Table 7.** Overview of the datasets that are used for testing the performance of the automated system. The table contains the number of the dataset from 1 to 5, the number of files in the dataset, the size of the dataset in MBs, the runtime in seconds for the 2017-method, and the runtime in seconds for the best performing method. Also, a total overview of the number of files, the size in MBs, the runtime in seconds for the 2017-method, and runtime for the best performing method.

| Dataset | Files | Size | Time (s) 2017-method | Time (s) Best method |
| --- | --- | --- | --- | --- |
| 1 | 80 | 202 MB | 100.6 | 59.2 |
| 2 | 147 | 231 MB | 171.3 | 94.1 |
| 3 | 281 | 605 MB | 440.8 | 367.6 |
| 4 | 549 | 1360 MB | 990.6 | 1049.5 |
| 5 | 1085 | 2960 MB | 2032.8 | 2176.8 |
| Total | 2142 | 5360 MB | 3736.1 | 3747.2 |

**Figure 7.**
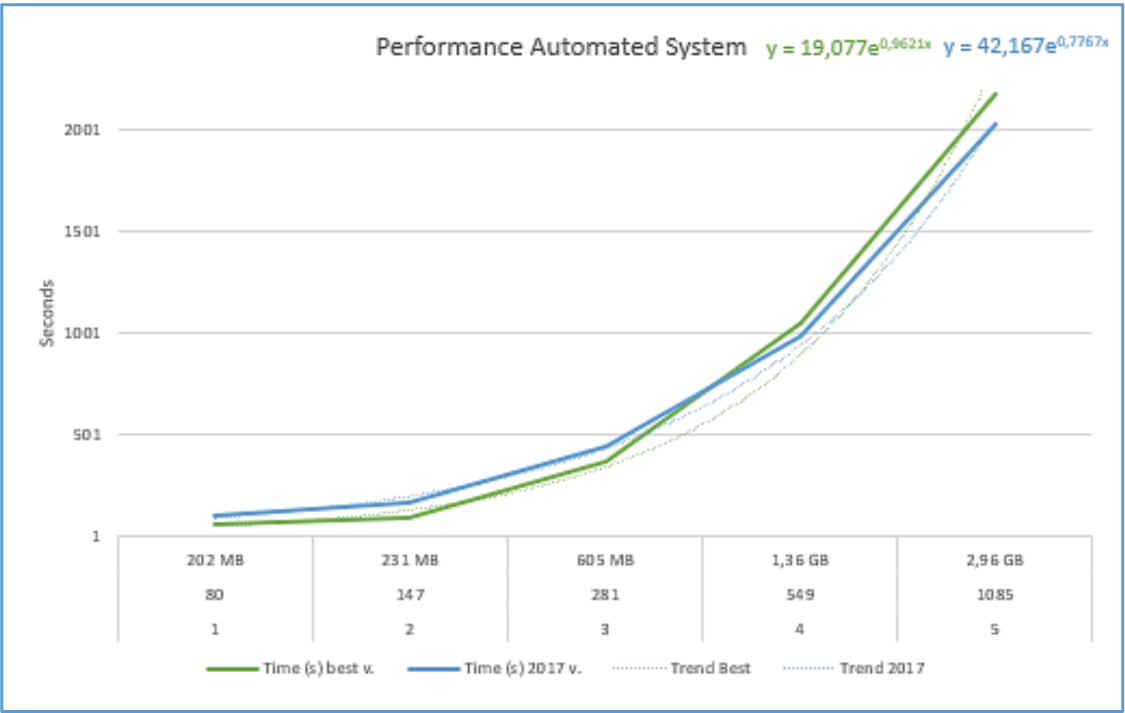
Graph showing the runtime, in seconds, needed to analyse the different sized datasets with the 2017-method and the best performing method. Trendlines are fitted to the lines and a formula is presented. The x-axis shows the number of files, size, and dataset number. The y-axis shows the runtime in seconds. The best performing method is coloured green, the 2017-method is coloured blue.

### Web application

At “<u>plakvallen.naturalis.nl</u>” the homepage of the web application can be found, as seen in figure 8. The web application allows users to upload, when navigating to the upload page, multiple photographs of sticky traps that were set in one meadow (maximum of 10 photographs). Users are asked to provide information about the meadow the traps were placed in and the traps itself. This includes the location and number of the meadow, date of trap placement, date of trap removal, height of the grass, mowing of the meadow, grazing of the meadow, measurement of sunshine, rain, and wind, quantity of biodiversity, type of fertilization (firm manure, rough manure, do not know), and management package (agriculture without management package, agriculture with postponed mowing, agriculture with extensive grazing, agriculture with nest management, agriculture with herb-rich grassland, nature with moist meadow, nature with meadow birds package, do not know). Information for a certain meadow and a setting date can only be transmitted as input once to avoid redundancy. The location of the meadow field can be chosen from a map that is provided with *Geoposition*. A part of the upload page is shown in figure 9.

**Figure 8.**
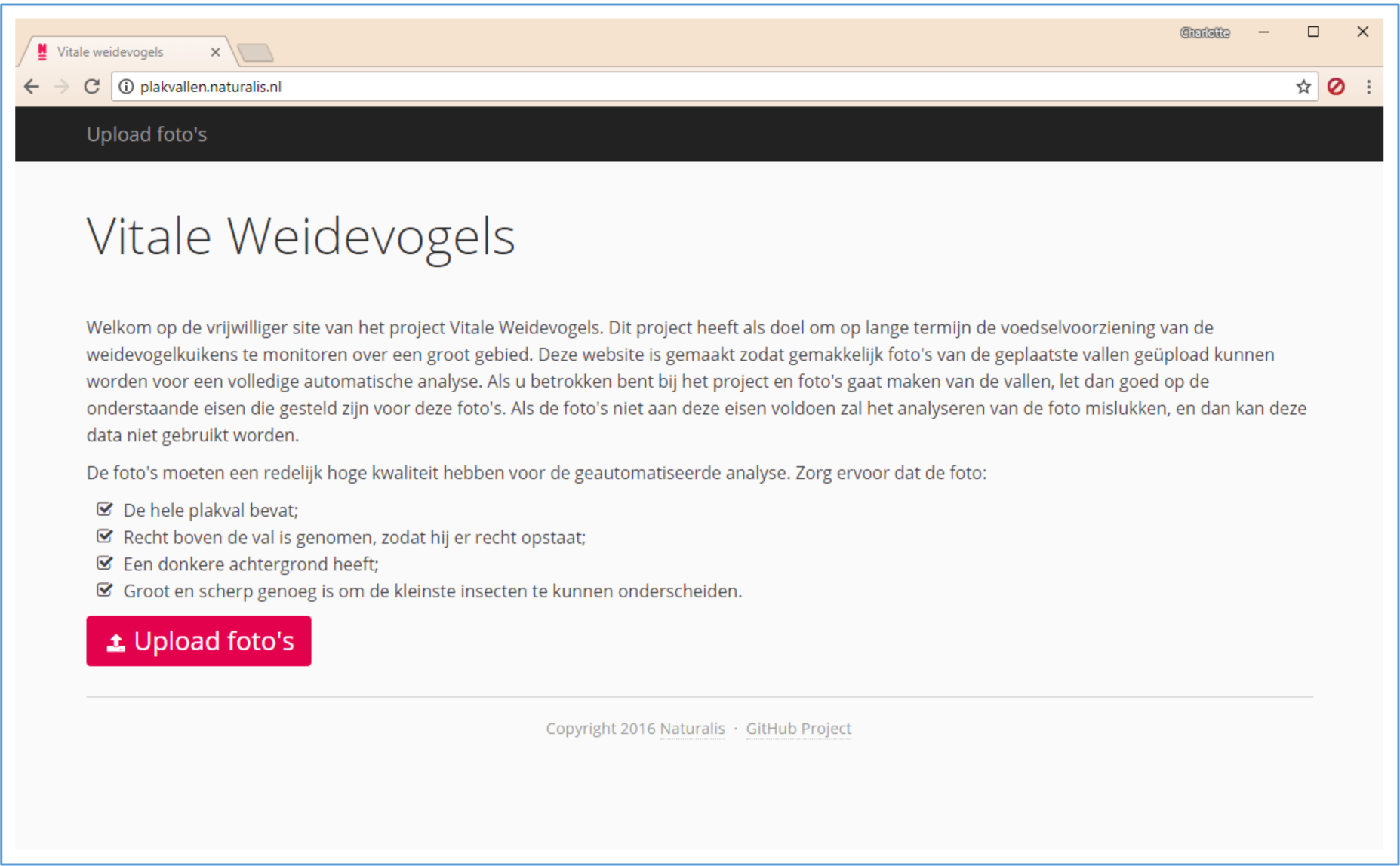
The homepage of plakvallen.naturalis.nl for the “Vitale Weidevogels” monitoring project. The homepage consists of an explanation of the project and the criteria for the photographs. Navigating to the upload page can be accomplished by clicking on “Upload foto’s” at the top of the page or the red button.

**Figure 9.**
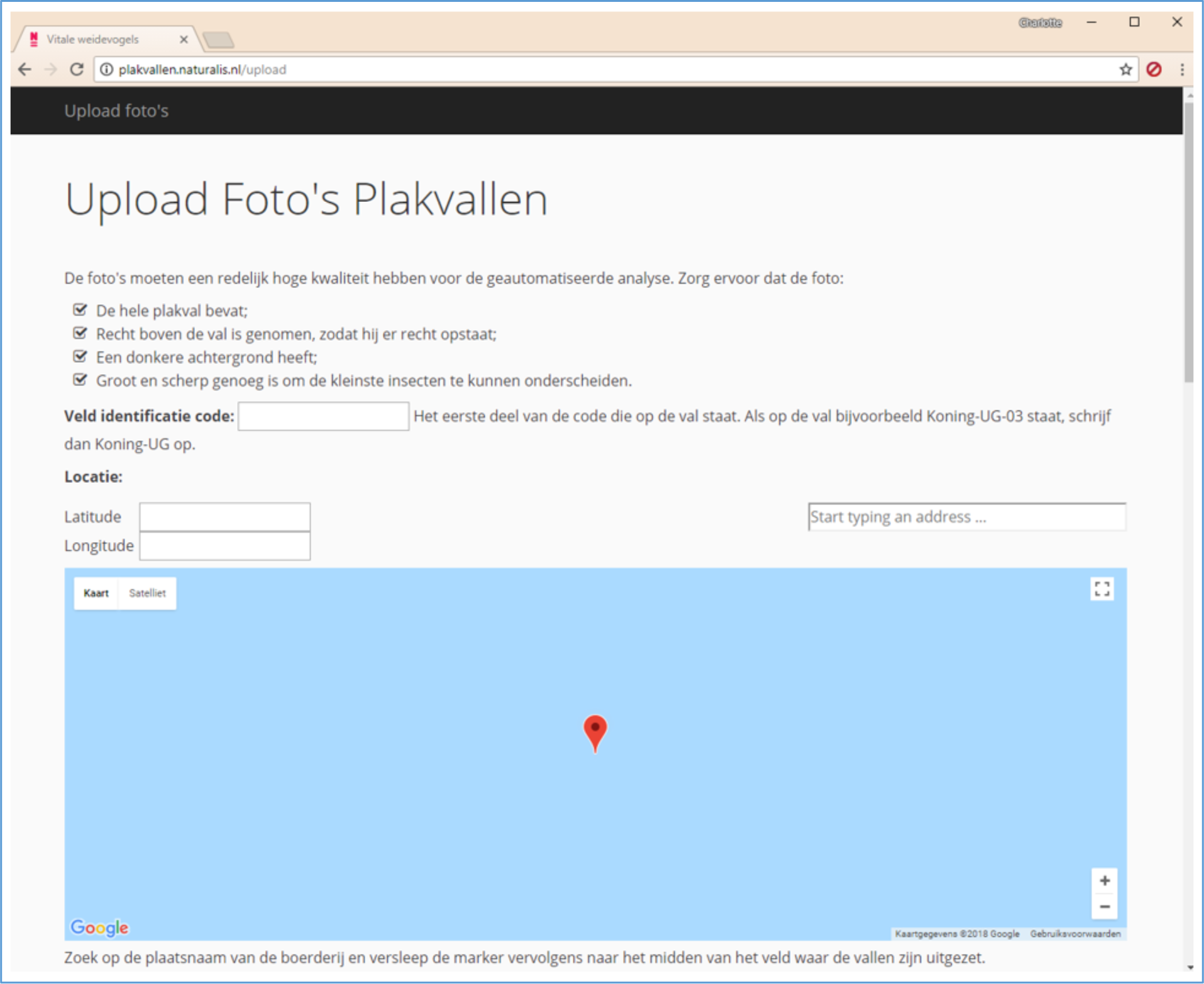
The top of the upload page of plakvallen.naturalis.nl for the “Vitale Weidevogels” monitoring project. On the upload page the criteria for the photographs are shown again, the field identification code can be entered and a location can be entered to select a meadow. The remainder of the information about the meadow and the traps can be specified below.

When photographs of traps have been uploaded and all images are analysed it will present the results for those images, as shown in figure 10. An average mm2 of insects per traps and mm2 variance is given. For each trap the total number of insects, the number of insects smaller than 4mm, the number of insects between 4 and 10mm, and the number of insects larger than 10mm is given. For each of those size classes an average is given. If an image cannot be analysed the web application still redirects to the result page and show results, however it does show a warning with a message about which trap number is the faulty image. The results, information and images are not saved in the database and the web application asks the user to upload the photographs of the traps again and to remake the photograph of the trap that has a faulty image. When no errors have occurred the images, information, and results are saved into the database.

**Figure 10.**
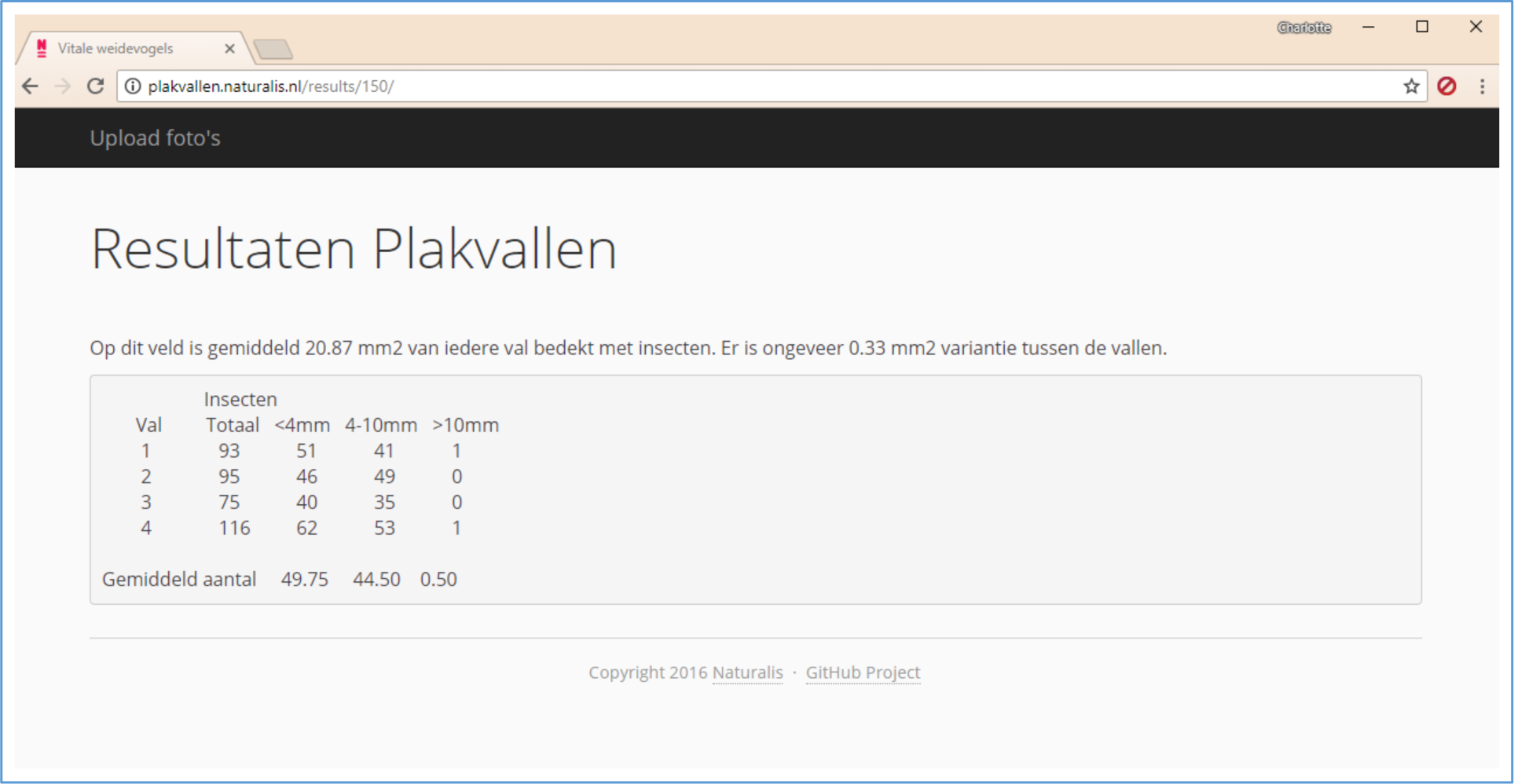
The result page of plakvallen.naturalis.nl for the “Vitale Weidevogels” monitoring project. On the result page the results for the uploaded images are presented. An average mm2 of insects for per traps and mm2 variance is given. For each trap the total number of insects, the number of insects smaller than 4mm, the number of insects between 4 and 10mm, and the number of insects larger than 10mm is given. For each of those classes an average is given. If an image cannot be analysed the web application still redirects to the result page and show results, however it does show a warning with a message about which trap number is the faulty image.

### Command line

Large numbers of traps are analysed in a fast and easy way with the command line version of *sticky-traps*. The results are printed inside the terminal and saved in a result file, if set in the *YAML* file, shown in figure 11. Figure 12 consists of a part of part of the *YAML* file that is used to specify preferences and information about the traps.

**Figure 11.**
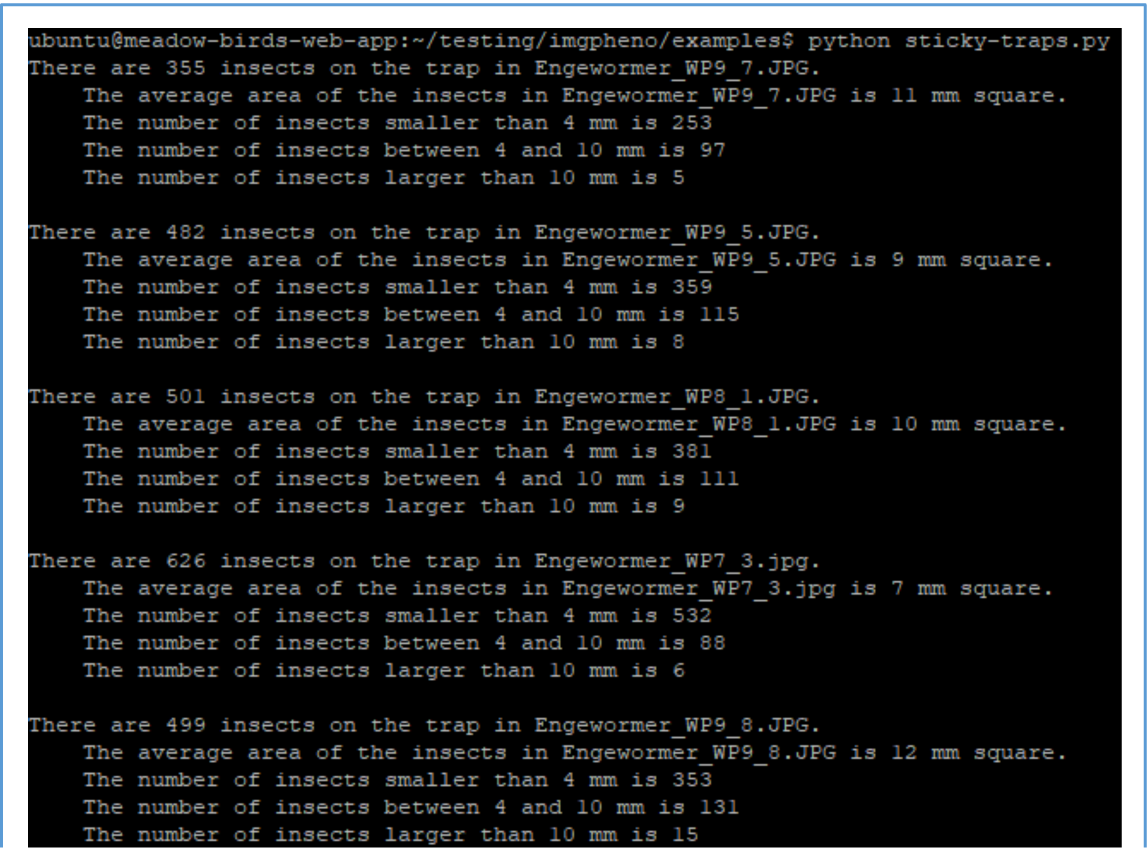
Screenshot of running sticky-traps command line program, in the command line terminal. The program is called with “python sticky-traps.py”, it prints the results for each of the analysed images in the terminal.

**Figure 12.**
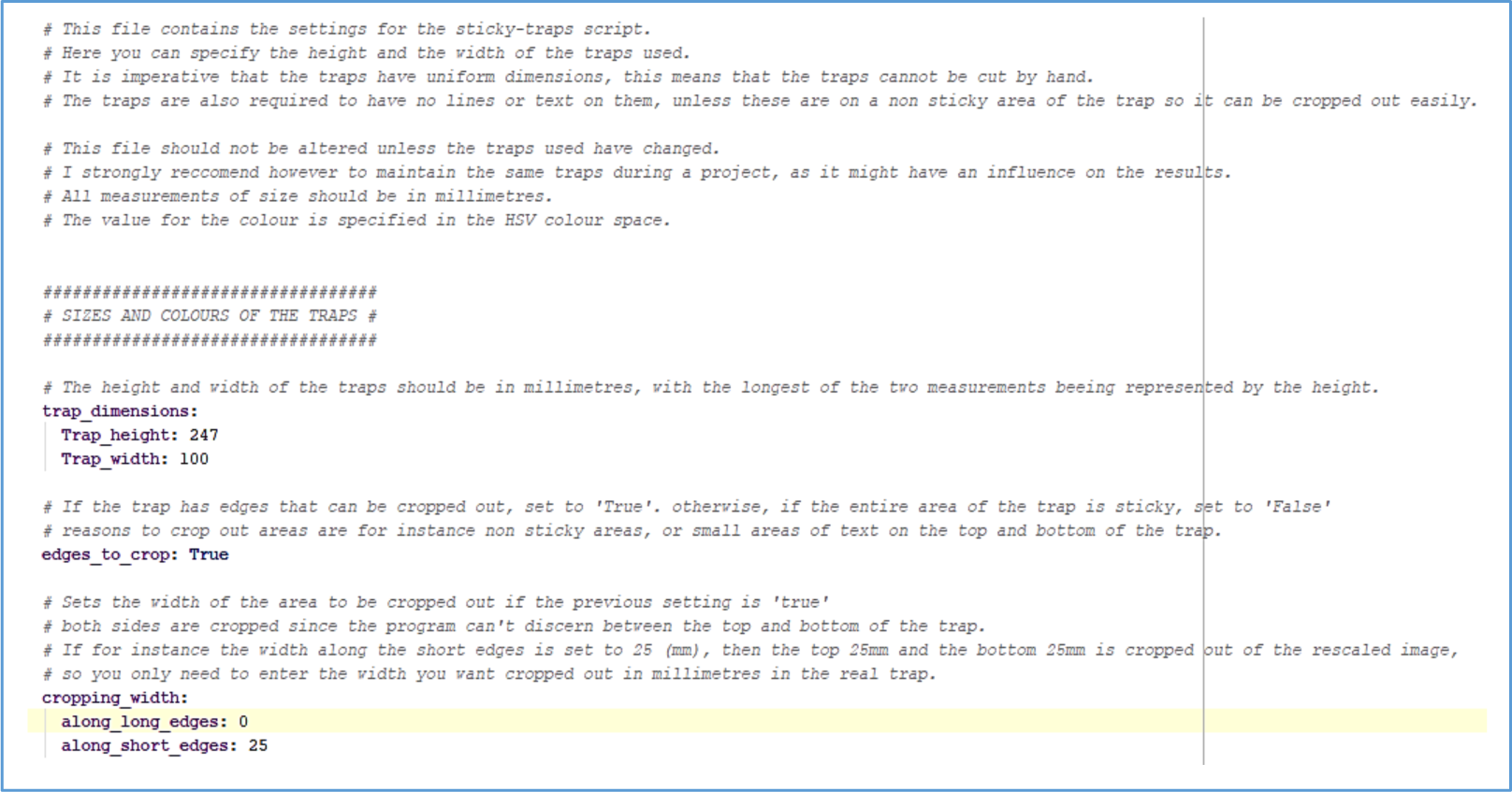
Top part of the YAML file “sticky-traps.yml”. The file consists of information about the settings that can be changed. In this part the dimensions, and cropping of the trap can be adjusted. Below those are the remainder of the settings.

### Analysis of the traps

All the traps, from both days in 2017 and 2018, are analysed with the best performing method (“Automatically counted 41, 22 - w/o. filter - changed cont. det. - w. limit”). Figure 13 shows a comparison of two traps; the left trap is a trap that was set on the 1^st^ day of 2017 and the right trap was set on the 1^st^ day of 2018. Ellipses are drawn, in red, for each contour that is found. The ellipses are not drawn well on image of the trap from 2017, this means that the contours too could not be found and drawn as reliable. Also, the perspective of the image of the trap from 2017 is not transformed correctly, causing the determination of the size of the insects to be inaccurate. The ellipses on the image of the trap from 2018 are drawn more accurately as this image is clearer and the trap is not in a protective plastic cover, therefore there are no air bubbles.

**Figure 13.**
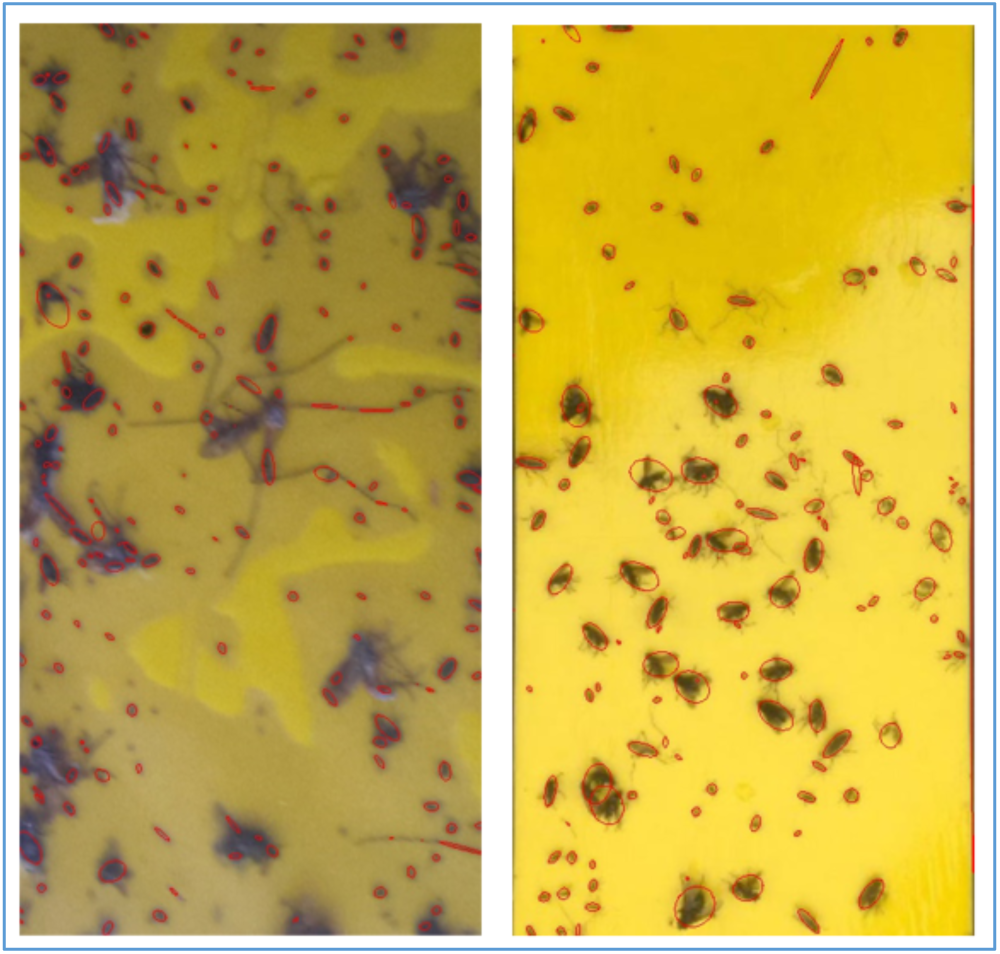
Drawn ellipses, in red, of the contours of insects on images from 2017 and 2018. The left image shows the ellipses drawn on a trap that was set on the 1^st^ day of 2017, on the right the same is done for an image of a trap that was set on the 1^st^ day of 2018.

The bell curves visualising distribution of the total insects that are counted on both days in 2017 and 2018, figure 14 and 15, show that the data is normally distributed. Averages for each meadow for each size class are calculated and two-sided paired t-tests are performed.

**Figure 14.**
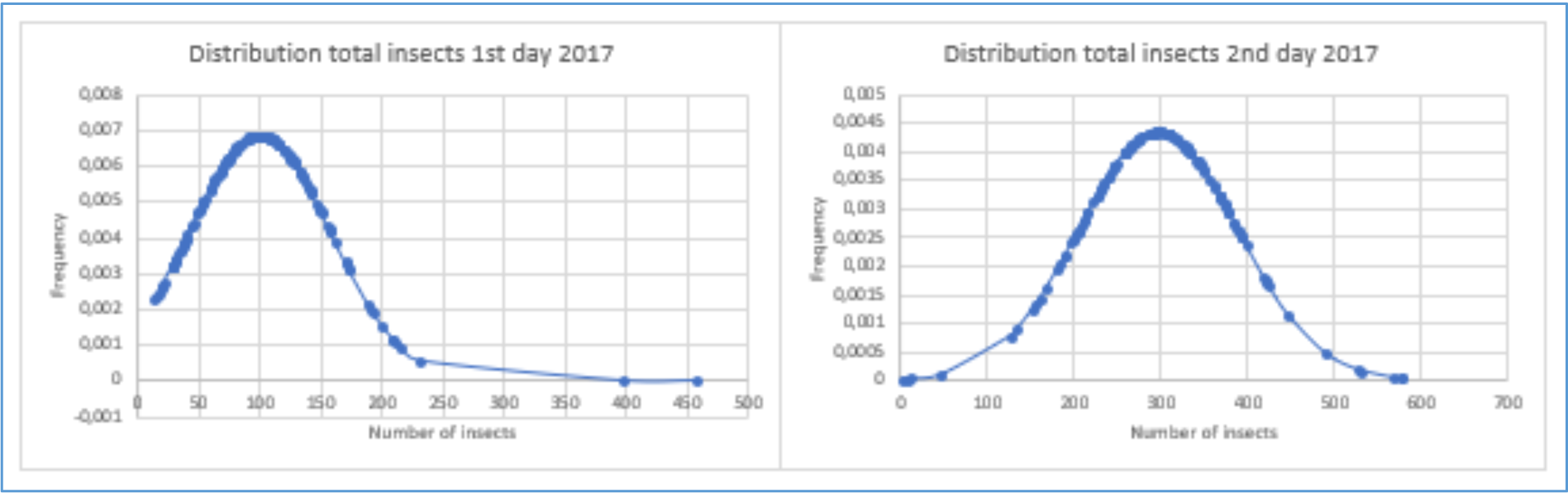
Histograms of the total insects counted on the 1^st^ and 2^nd^ day of 2017. On the left the distribution is visualised for the 1^st^ day of 2017, and on the right the 2^nd^ day is visualised. The x-axis represents the number of insects and the y-axis represents the frequency.

**Figure 15.**
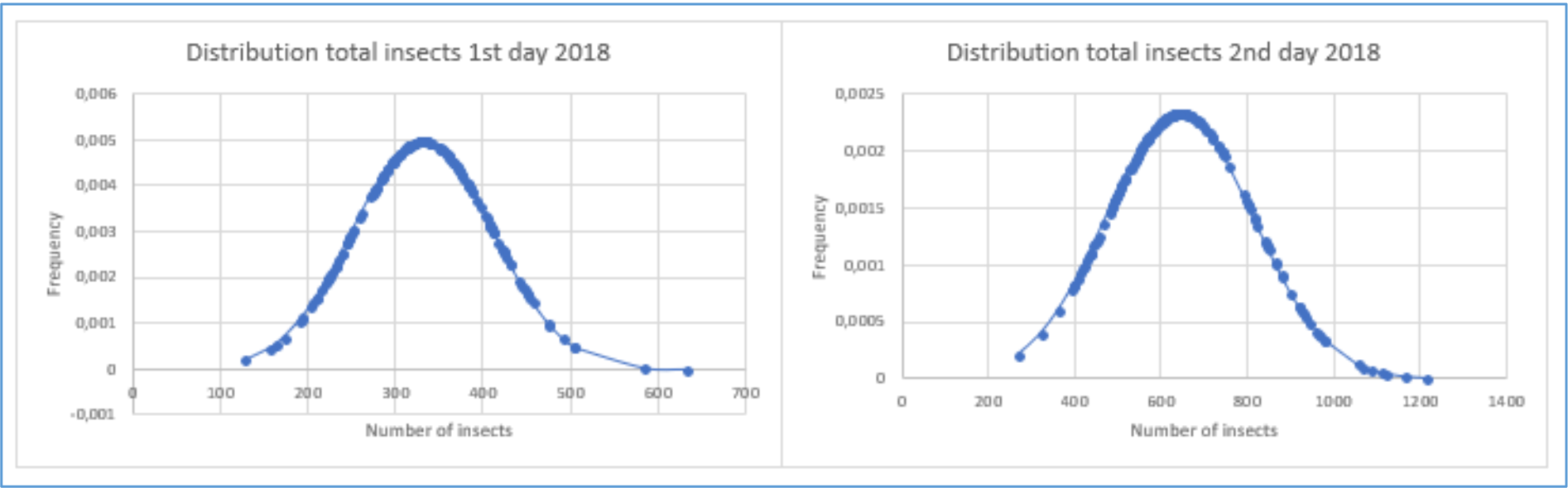
Histograms of the total insects counted on the 1^st^ and 2^nd^ day of 2018. On the left the distribution is visualised for the 1^st^ day of 2018, and on the right the 2^nd^ day is visualised. The x-axis represents the number of insects and the y-axis represents the frequency.

The results of the performed t-tests indicate that for each class there is significant difference between 2017 and 2018, as seen in table 8. Some values are removed (Koning-UM, Westerneng-LB) due to missing values, photographs of traps, in 2017. This would mean that the number of insects in 2018 are significantly greater when looking at the total averages.

**Table 8.** The results of the two-sided paired t-tests comparing all size classes of 2017 with 2018 for both days. The table holds the p-value for each test and the total averages of 2017 and 2018 for the tested size classes for each day. Significant differences (α = 0.05) indicated by asterisks *.

| Category | *p*-value | Average 2017 | Average 2018 |
| --- | --- | --- | --- |
| Total 2017 vs. 2018 1<sup>st</sup> day | 4.166 x 10<sup>-12</sup> * | 99.124 | 332.385 |
| Total 2017 vs. 2018 2<sup>nd</sup> day | 3.321 x 10<sup>-08</sup> * | 286.745 | 648.488 |
| <4mm 2017 vs. 2018 1<sup>st</sup> day | 6.476 x 10<sup>-13</sup> * | 60.482 | 227.991 |
| <4mm 2017 vs. 2018 2<sup>nd</sup> day | 2.938 x 10<sup>-08</sup> * | 215.522 | 517.088 |
| 4-10mm 2017 vs. 2018 1<sup>st</sup> day | 3.859 x 10<sup>-09</sup> * | 36.531 | 97.078 |
| 4-10mm 2017 vs. 2018 2<sup>nd</sup> day | 1.381x10<sup>-04</sup> * | 75.712 | 125.608 |
| >10mm 2017 vs. 2018 1<sup>st</sup> day | 7.192x10<sup>-05</sup> * | 2.377 | 6.153 |
| >10mm 2017 vs. 2018 2<sup>nd</sup> day | 0.022 | 4.201 | 6.231 |

The results of the two-sided t-tests, in table 9, with equal or unequal variances (depending on the results of the f-tests, which are not included) to determine if the difference for each size class for each day per meadow in 2017 versus 2018 is significant show that most are significant. Except for Engewormer-WP5 on the 2^nd^ day in the 4-10mm size class, and Engewormer-VH2, Engewormer-VH3, Engewormer-WP1, Engewormer-WP5, Engewormer-WP6, Hudding-KG, Hudding-LB, and Westerneng-UM on the 2^nd^ day in the >10mm class.

**Table 9.**
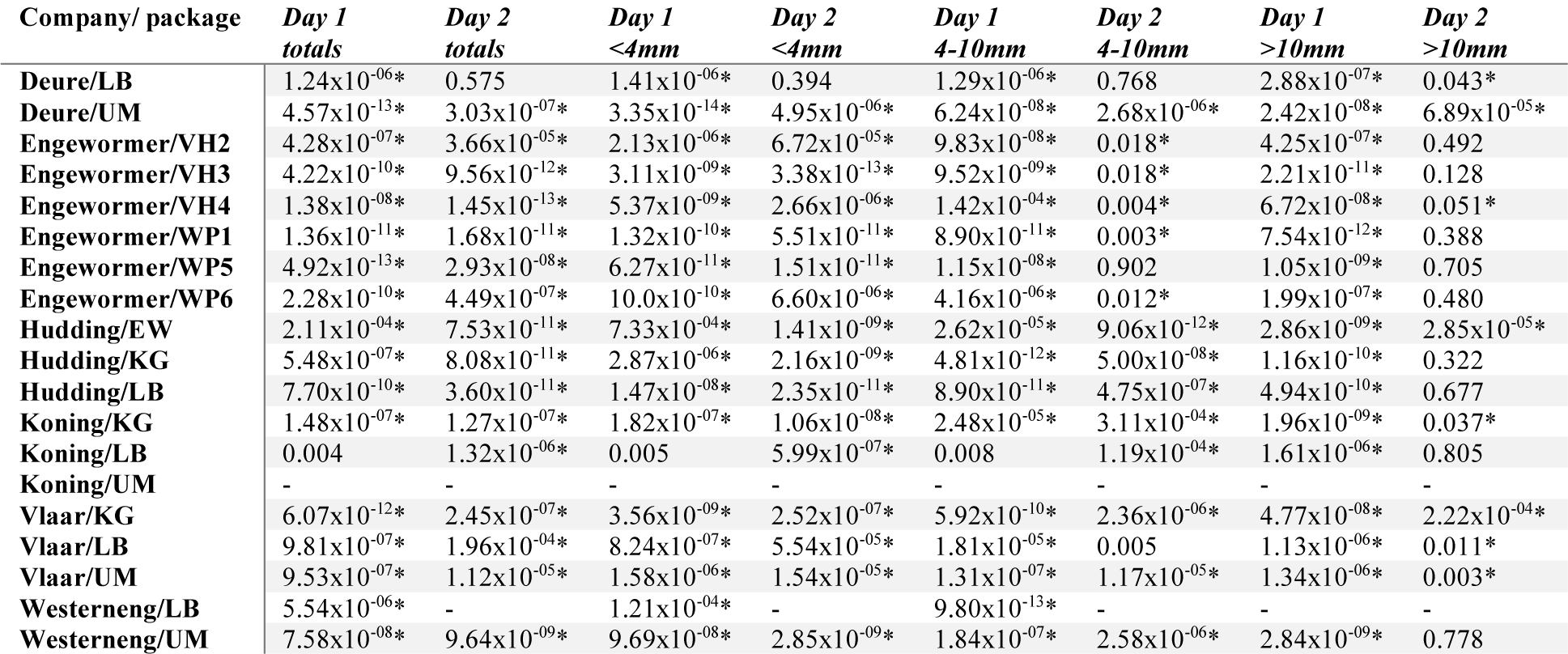
Results of two-sided t-tests comparing insect counts for each day, for each meadow (i.e. company/management combination) between 2017 and 2018. Significant differences (α = 0.05) indicated by asterisks *. The entries with ‘-’ mean that the data for the meadow was missing completely for the year 2017.

| Company/ package | Day 1<br>totals | Day 2<br>totals | Day 1<br><4mm | Day 2<br><4mm | Day 1<br>4-10mm | Day 2<br>4-10mm | Day 1<br>>10mm | Day 2<br>>10mm |
| --- | --- | --- | --- | --- | --- | --- | --- | --- |
| Deure/LB | 1.24x10<sup>-06</sup>* | 0.575 | 1.41x10<sup>-06</sup>* | 0.394 | 1.29x10<sup>-06</sup>* | 0.768 | 2.88x10<sup>-07</sup>* | 0.043* |
| Deure/UM | 4.57x10<sup>-13</sup>* | 3.03x10<sup>-07</sup>* | 3.35x10<sup>-14</sup>* | 4.95x10<sup>-06</sup>* | 6.24x10<sup>-08</sup>* | 2.68x10<sup>-06</sup>* | 2.42x10<sup>-08</sup>* | 6.89x10<sup>-05</sup>* |
| Engewormer/VH2 | 4.28x10<sup>-07</sup>* | 3.66x10<sup>-05</sup>* | 2.13x10<sup>-06</sup>* | 6.72x10<sup>-05</sup>* | 9.83x10<sup>-08</sup>* | 0.018* | 4.25x10<sup>-07</sup>* | 0.492 |
| Engewormer/VH3 | 4.22x10<sup>-10</sup>* | 9.56x10<sup>-12</sup>* | 3.11x10<sup>-09</sup>* | 3.38x10<sup>-13</sup>* | 9.52x10<sup>-09</sup>* | 0.018* | 2.21x10<sup>-11</sup>* | 0.128 |
| Engewormer/VH4 | 1.38x10<sup>-08</sup>* | 1.45x10<sup>-13</sup>* | 5.37x10<sup>-09</sup>* | 2.66x10<sup>-06</sup>* | 1.42x10<sup>-04</sup>* | 0.004* | 6.72x10<sup>-08</sup>* | 0.051* |
| Engewormer/WP1 | 1.36x10<sup>-11</sup>* | 1.68x10<sup>-11</sup>* | 1.32x10<sup>-10</sup>* | 5.51x10<sup>-11</sup>* | 8.90x10<sup>-11</sup>* | 0.003* | 7.54x10<sup>-12</sup>* | 0.388 |
| Engewormer/WP5 | 4.92x10<sup>-13</sup>* | 2.93x10<sup>-08</sup>* | 6.27x10<sup>-11</sup>* | 1.51x10<sup>-11</sup>* | 1.15x10<sup>-08</sup>* | 0.902 | 1.05x10<sup>-09</sup>* | 0.705 |
| Engewormer/WP6 | 2.28x10<sup>-10</sup>* | 4.49x10<sup>-07</sup>* | 10.0x10<sup>-10</sup>* | 6.60x10<sup>-06</sup>* | 4.16x10<sup>-06</sup>* | 0.012* | 1.99x10<sup>-07</sup>* | 0.480 |
| Hudding/EW | 2.11x10<sup>-04</sup>* | 7.53x10<sup>-11</sup>* | 7.33x10<sup>-04</sup>* | 1.41x10<sup>-09</sup>* | 2.62x10<sup>-05</sup>* | 9.06x10<sup>-12</sup>* | 2.86x10<sup>-09</sup>* | 2.85x10<sup>-05</sup>* |
| Hudding/KG | 5.48x10<sup>-07</sup>* | 8.08x10<sup>-11</sup>* | 2.87x10<sup>-06</sup>* | 2.16x10<sup>-09</sup>* | 4.81x10<sup>-12</sup>* | 5.00x10<sup>-08</sup>* | 1.16x10<sup>-10</sup>* | 0.322 |
| Hudding/LB | 7.70x10<sup>-10</sup>* | 3.60x10<sup>-11</sup>* | 1.47x10<sup>-08</sup>* | 2.35x10<sup>-11</sup>* | 8.90x10<sup>-11</sup>* | 4.75x10<sup>-07</sup>* | 4.94x10<sup>-10</sup>* | 0.677 |
| Koning/KG | 1.48x10<sup>-07</sup>* | 1.27x10<sup>-07</sup>* | 1.82x10<sup>-07</sup>* | 1.06x10<sup>-08</sup>* | 2.48x10<sup>-05</sup>* | 3.11x10<sup>-04</sup>* | 1.96x10<sup>-09</sup>* | 0.037* |
| Koning/LB | 0.004 | 1.32x10<sup>-06</sup>* | 0.005 | 5.99x10<sup>-07</sup>* | 0.008 | 1.19x10<sup>-04</sup>* | 1.61x10<sup>-06</sup>* | 0.805 |
| Koning/UM | - | - | - | - | - | - | - | - |
| Vlaar/KG | 6.07x10<sup>-12</sup>* | 2.45x10<sup>-07</sup>* | 3.56x10<sup>-09</sup>* | 2.52x10<sup>-07</sup>* | 5.92x10<sup>-10</sup>* | 2.36x10<sup>-06</sup>* | 4.77x10<sup>-08</sup>* | 2.22x10<sup>-04</sup>* |
| Vlaar/LB | 9.81x10<sup>-07</sup>* | 1.96x10<sup>-04</sup>* | 8.24x10<sup>-07</sup>* | 5.54x10<sup>-05</sup>* | 1.81x10<sup>-05</sup>* | 0.005 | 1.13x10<sup>-06</sup>* | 0.011* |
| Vlaar/UM | 9.53x10<sup>-07</sup>* | 1.12x10<sup>-05</sup>* | 1.58x10<sup>-06</sup>* | 1.54x10<sup>-05</sup>* | 1.31x10<sup>-07</sup>* | 1.17x10<sup>-05</sup>* | 1.34x10<sup>-06</sup>* | 0.003* |
| Westerneng/LB | 5.54x10<sup>-06</sup>* | - | 1.21x10<sup>-04</sup>* | - | 9.80x10<sup>-13</sup>* | - | - | - |
| Westerneng/UM | 7.58x10<sup>-08</sup>* | 9.64x10<sup>-09</sup>* | 9.69x10<sup>-08</sup>* | 2.85x10<sup>-09</sup>* | 1.84x10<sup>-07</sup>* | 2.58x10<sup>-06</sup>* | 2.84x10<sup>-09</sup>* | 0.778 |

The p-value, of 0.841, for the single factor ANOVA that is performed on the grouped, per management package, total insect count averages for each meadow on both days in 2017 shows that there is no significant difference in insect counts between management packages, and the p-value, of 0.999, for the single factor ANOVA that is performed on the grouped, per management package, total insect count averages for each meadow on both days in 2018 shows that there is no significant difference in insect counts between management packages.

## DISCUSSION

As not all of the traps have been counted by hand and only hand counted results of traps from 2017 are available the reliability of the automated system is difficult to determine and to improve. However, the current automated system (using the best performing method for the algorithm) does show a lot of improvement of accuracy, as compared to the 2017-method, mainly for the vol images. This improvement of accuracy is even seen in bad quality, blurry/poor lighting/in protective plastic cover, photographs of traps. The current automated system is focussed on good quality photographs of traps without a protective plastic cover, so it is assumed that the results should be more accurate when used to analyse good quality photographs.

Although improvement of accuracy is visible it will be difficult to improve the accuracy even further unless there is a standardised way of making photographs of traps. For both methods of the algorithm there are only two categories, 4-10 mm 1^st^ day and >10mm 2^nd^ day for the 2017-method, and 4-10mm 2^nd^ day and >10mm 1^st^ day for the best performing method, in which the difference between the prof and vol images is insignificant, so the results are heavily influenced by the quality of the photograph and two different photographs of the same trap do not mean that the results are insignificantly different, as seen in table 6.

The photographs of the traps that are taken in 2018 all are of very good quality and if this level of quality is maintained the accuracy is easier to improve and the change in the number of insects will be easier to determine. As the photographs of the traps from 2017 overall are quite poor quality it is questioned how reliable the results are and how reliable the conclusions are that can be drawn from these results. Not all photographs of the traps were available or some were unable to be analysed, this leaves some gaps in the results and again reduces the reliability.

Despite the aforementioned issues, the current automated system does show a large improvement as compared to the automated system using the 2017-method. For only one of the categories, 4-10mm 2^nd^ day, for both prof and vol images the t-test resulted in an insignificant difference between the hand counted results and the automatically counted results using the 2017-method. For two of the categories, total 2^nd^ day and <4mm 2^nd^ day, for both prof and vol images the t-tests resulted in an insignificant difference, and for three of the categories, total 1^st^ day vol, <4mm 1^st^ day vol and 4-10mm 1^st^ day prof, the t-tests resulted in an insignificant difference as well, as seen in table 5. This means that the results of the current automated system are more equal to the hand counted results and are likely to be more reliable.

The current automated system is faster when using a smaller dataset. This version of the automated system is thus a better option for the web application as it only analyses 10 images at a time, shown in table 7. The 2017-method performs a little better when analysing larger datasets, however the total runtime of the analysis of all the datasets only differ 11.1 seconds between the 2017-method and the best performing method.

For some of the volunteers from ANV Water, Land & Dijken, who were the users of the web application, uploading the images went without any issues and mentioned the web application was easy to use, and for some of the volunteers uploading the photographs and using the web application was difficult. Most of the volunteers do not have ample experience with computers and using the internet, as these are key components for using the web application this does not aid in user friendly uploading of the photographs and therefore does not fulfil its purpose. The web application will need further adjustment to provide user friendly uploading for all of its users. User feedback on the result page was overwhelmingly positive, instantly seeing results of the traps was very welcome.

As mentioned before, it is questioned how reliable the results of the comparison of the traps that were set in 2017 and the traps that were set in 2018 are because the quality of the images is very diverse and some images are missing or could not be analysed. The results from the t-tests, table 8, indicate that there is significant difference between the number of insects in 2017 versus 2018 for all size classes and days. The averages of counted insects for 2018 are a lot higher as compared to the averages of counted insects for 2017, thus the number of insects is significantly higher in 2018.

The average counted insects for 2018 are in fact so much higher that the authenticity of the results is questioned. In one year an insect population could have grown this much, as insect growth is exponential, however it does greatly depend on the weather (Birch, 1948). When the traps were set in 2018 the temperature on the 1^st^ day was 25 °C at maximum and 29 °C at maximum on the 2^nd^ day (KNMI, 2018b), in 2017 the temperature on the 1^st^ day was 13 °C at maximum and 21 °C at maximum on the 2^nd^ day (KNMI, 2018a). This difference in temperature could indicate why so much more insects have been counted in 2018, as this would not necessarily mean that the population has grown, but that the high temperatures caused more insects to have metamorphosed from larvae into adult flying insects earlier than in 2017.

In warmer temperatures insects grow faster but they produce smaller body sizes, in colder temperatures growth rate is slower but insect bodies are larger (Chen, Leask, MacKinnon, Ramanaden, & Yoon, 2014). As seen in table 5, in 2017 1.7% of all insects that are detected are larger than 10mm, in 2017 1.3% of all insects that are detected are larger than 10mm, this could indicate that the high temperatures caused more insects to have grown to adult size but less insects have grown larger than 10mm. Another possible explanation for the large increase in insects is that the traps in 2017 were less sticky than in 2018 and caused more insects to escape the traps. Yet, the same traps were used in 2017 and 2018, and assuming the traps that were set in 2018 would have lost their degree of stickiness would be more probable as the traps were one year old. Most likely, the poor quality of the images in 2017 could have led to the inability to find all the contours of the insects on the trap and resulted in fewer insects.

In table 9 the results of the t-tests show that in almost all meadows insect numbers are significantly increased, except for Deure-LB (total 2^nd^ day, <4mm 2^nd^ day, 4-10mm 2^nd^ day), Engewormer-WP5 (4-10mm 2^nd^ day, >10mm 2^nd^ day), and Engewormer-VH2, Engewormer-VH3, Engewormer-WP1, Engewormer-WP6, Hudding-KG, Hudding-LB, Koning-LB and Westerneng-UM (all >10mm 2^nd^ day). This could indicate that most of the management packages have a positive influence on insect population growth. For both 2017 and 2018 there seems to be no significant difference in insect numbers between management packages in the same year and this reinforces the indication that most management packages have a positive influence on insect population growth.

For any follow-up research it is imperative that the photographs of the traps at least continue consisting of a high quality. It would also be more valuable to set traps in more meadows and to set them on more than two days to gain a better overview of the insect counts in different types of weather.

## CONCLUSIONS

To answer if the food supply for meadow birds in the Netherlands is sufficient, the reliability of the results has to be taken into consideration. If the results are accurate then the number of insects found on sticky traps that were deployed in the Netherlands has increased and no specific correlation can be found between insect number and the provided management package. The results indicate that each of the management packages has a positive influence on insect population growth. If an average of 300 up to 650 insects on a single trap in 2018 are found, as compared to an average of 99 up to 290 insects on a single trap in 2017, then this could imply that the food supply for meadow birds is at least more sufficient than it was in 2017. If the insect populations have grown as significantly as is indicated from the results then it would be probable that an increase in meadow birds has occurred or will occur in the near future.

## SUPPLEMENTARY MATERIALS

- The source code for the web application can be found on GitHub: https://github.com/naturalis/nbclassify/tree/sticky-traps/html/sticky_traps

## LITERATURE

ANV Water Land & Dijken. (2018). Photograph of deployed sticky trap. Retrieved from https://twitter.com/waterlanddijken/status/993814485969395714

Beintema, A. J., Moedt, O., & Ellinger, D. (1995). Ecologische Atlas van de Nederlandse weidevogels.

Beintema, A. J., & Visser, G. H. (1989). The effect on weather on time budgets and development of chicks of meadow birds. Ardea, 77(2), 181–192. Retrieved from http://ardea.nou.nu/ardeapdf//a77-181-192.pdf

Ben-Kiki, O., & Evans, C. (2001). YAML Ain’t Markup Language (YAML™) Version 1.2 YAML Ain’t Markup Language (YAML™) Version 1.2 3 rd Edition, Patched at 2009-10-01. Retrieved from http://yaml.org/spec/1.2/spec.html

Benton, T. G., Bryant, D. M., Cole, L., & Crick, H. Q. P. (2002). Linking agricultural practice to insect and bird populations: a historical study over three decades. Journal of Applied Ecology, 39(4), 673–687. https://doi.org/10.1046/j.1365-2664.2002.00745.x

Birch, L. C. (1948). The Intrinsic Rate of Natural Increase of an Insect Population. Journal of Animal Ecology, 17(1), 15–26. Retrieved from https://www.britishecologicalsociety.org/100papers/100_Ecological_Papers/100_Influential_Papers_003.pdf

Boele, A., van Bruggen, J., Hustings, F., Koffijberg, K., Vergeer, J.-W., van der Meij met medewerking van Symen Deuzeman, T., … van der Jeugd, H. (2016). Broedvogels in Nederland in 2014. Retrieved from https://www.sovon.nl/sites/default/files/doc/Rap_2016-04_Broedvogelrapport-2014-LR.pdf

Breeuwer, A., Berendse, F., Willems, F., Foppen, R., Teunissen, W., Schekkerman, H., & Goedhart, P. (2009). Do meadow birds profit from agri-environment schemes in Dutch agricultural landscapes? Biological Conservation, 142(12), 2949–2953. https://doi.org/10.1016/J.BIOCON.2009.07.020

CBS, & AgroXpertus. (2018). CBS StatLine - Grasland; oppervlakte en opbrengst. Retrieved February 28, 2018, from http://statline.cbs.nl/StatWeb/publication/?DM=SLNL&PA=7140gras&D1=0-1&D2=a&D3=15,20,25-26,l&VW=T

Chamberlain, D. E., Fuller, R. J., Bunce, R. G. H., Duckworth, J. C., & Shrubb, M. (2000). Changes in the abundance of farmland birds in relation to the timing of agricultural intensification in England and Wales. Journal of Applied Ecology, 37(5), 771–788. https://doi.org/10.1046/j.1365-2664.2000.00548.x

Chen, S. Y., Leask, K. P., MacKinnon, S. W., Ramanaden, Y. J., & Yoon, J. H. (2014). The effects of temperature on the time to maturation of Drosophila melanogaster. The Expedition, 3(0). Retrieved from http://ojs.library.ubc.ca/index.php/expedition/article/view/184797

Donald, P. F., Green, R. E., & Heath, M. F. (2001). Agricultural intensification and the collapse of Europe’s farmland bird populations. Proceedings. Biological Sciences, 268(1462), 25–29. https://doi.org/10.1098/rspb.2000.1325

Donald, P. F., Sanderson, F. J., Burfield, I. J., & van Bommel, F. P. J. (2006). Further evidence of continent-wide impacts of agricultural intensification on European farmland birds, 1990–2000. Agriculture, Ecosystems & Environment, 116(3–4), 189–196. https://doi.org/10.1016/J.AGEE.2006.02.007

ETI Bioinformatics. (2018). SoortenBank.nl: Vogels: Woordenlijst: voeden. Retrieved February 28, 2018, from http://www.soortenbank.nl/soorten.php?soortengroep=vogels&selected=definitie&menuentry=woordenlijst&record=voeden

Gamero, A., Brotons, L., Brunner, A., Foppen, R., Fornasari, L., Gregory, R. D., … Voříšek, P. (2017). Tracking Progress Toward EU Biodiversity Strategy Targets: EU Policy Effects in Preserving its Common Farmland Birds. Conservation Letters, 10(4), 395–402. https://doi.org/10.1111/conl.12292

Gregory, R. D., van Strien, A., Vorisek, P., Gmelig Meyling, A. W., Noble, D. G., Foppen, R. P. B., & Gibbons, D. W. (2005). Developing indicators for European birds. Philosophical Transactions of the Royal Society B: Biological Sciences, 360(1454), 269–288. https://doi.org/10.1098/rstb.2004.1602

Hallmann, C. A., Sorg, M., Jongejans, E., Siepel, H., Hofland, N., Schwan, H., … de Kroon, H. (2017). More than 75 percent decline over 27 years in total flying insect biomass in protected areas. PLOS ONE, 12(10), e0185809. https://doi.org/10.1371/journal.pone.0185809

KNMI. (2018a). Weerstatistieken KNMI - Weergegevens De Bilt Mei 2017. Retrieved June 20, 2018, from https://weerstatistieken.nl/de-bilt/2017/mei

KNMI. (2018b). Weerstatistieken KNMI - Weergegevens De Bilt Mei 2018. Retrieved June 20, 2018, from https://weerstatistieken.nl/de-bilt/2018/mei

Louis Bolk Instituut. (2015). Geen bescherming zonder voedsel. Retrieved from http://edepot.wur.nl/353511

Michels, R. (2017). Automating sticky trap analysis. Retrieved from https://github.com/naturalis/nbclassify-data/tree/master/archive-sticky-traps

Ministerie van Landbouw Natuur en Voedselkwaliteit. (2018). Rode lijsten: soort van Rode Lijst Vogels Beschermde natuur in Nederland. Retrieved February 28, 2018, from http://minez.nederlandsesoorten.nl/content/rode-lijsten-soort-van-rode-lijst-vogels

Naturalis Biodiversity Center. (2017). NBClassify. Retrieved from https://github.com/naturalis/nbclassify

Naturalis Biodiversity Center. (2018). naturalis/imgpheno: Image feature extraction in Python. Retrieved February 28, 2018, from https://github.com/naturalis/imgpheno

Pereira, S., Gravendeel, B., Wijntjes, P., & Vos, R. (2016). OrchID: a Generalized Framework for Taxonomic Classification of Images Using Evolved Artificial Neural Networks. BioRxiv, 070904. https://doi.org/10.1101/070904

Robinson, R. A., & Sutherland, W. J. (2002). Post-war changes in arable farming and biodiversity in Great Britain. Journal of Applied Ecology, 39(1), 157–176. https://doi.org/10.1046/j.1365-2664.2002.00695.x

Schekkerman, H., Teunissen, W., & Oosterveld, E. (2009). Mortality of Black-tailed Godwit Limosa limosa and Northern Lapwing Vanellus vanellus chicks in wet grasslands: influence of predation and agriculture. Journal of Ornithology, 150(1), 133–145. https://doi.org/10.1007/s10336-008-0328-4

Sovon. (2012). Factsheet aantallen boerenlandvogels over de laatste 50 jaar | Sovon.nl. Retrieved February 28, 2018, from www.sovon.nl/nl/content/factsheet-aantallen-boerenlandvogels-over-de-laatste-50-jaar

Teunissen, W., Schekkerman, H., Willems, F., & Majoor, F. (2008). Identifying predators of eggs and chicks of Lapwing Vanellus vanellus and Black-tailed Godwit Limosa limosa in the Netherlands and the importance of predation on wader reproductive output. Ibis, 150(1), 74–85. https://doi.org/10.1111/j.1474-919X.2008.00861.x

van Noordwijk, A. J., & Thomson, D. L. (2008). Survival Rates of Black-Tailed Godwits *Limosa limosa* Breeding in the Netherlands Estimated from Ring Recoveries. Ardea, 96(1), 47–57. https://doi.org/10.5253/078.096.0106

